# Extrastriate Connectivity of the Mouse Dorsal Lateral Geniculate Thalamus

**DOI:** 10.1101/351528

**Authors:** Michael S. Bienkowski, Nora L. Benavidez, Kevin Wu, Lin Gou, Marlene Becerra, Hong-Wei Dong

## Abstract

The mammalian visual system is one of the most well-studied brain systems. Visual information from the retina is relayed to the dorsal lateral geniculate nucleus of the thalamus (LGd). The LGd then projects topographically to primary visual cortex (VISp) to mediate visual perception. In this view, the VISp is a critical network hub where visual information must traverse LGd-VISp circuits to reach higher-order ‘extrastriate’ visual cortices. However, decades of conflicting reports in a variety of mammals support or refute the existence of extrastriate LGd connections that can bypass the VISp. Here, we provide evidence of bidirectional extrastriate connectivity with the mouse LGd. Using small, discrete coinjections of anterograde and retrograde tracers within the thalamus and cortex, our cross-validated approach identified bidirectional thalamocortical connectivity between LGd and extrastriate visual cortices. Our findings support the existence of extrastriate LGd circuits and provide novel understanding of LGd organization in rodent visual system.

---

In all mammals, visual information from the retina is projected topographically onto the dorsal lateral geniculate nucleus of the thalamus (LGd), which in turn projects a retinotopic map onto the primary visual cortex (VISp, also known as V1 or striate cortex). The VISp is positioned as a visual gateway to the rest of the cortex and this view is supported by the loss of vision caused by VISp damage. However, reports of visually-dependent behavior and visually-evoked activation of extrastriate cortex in cortically-blind patients has suggested additional neural circuit pathways that convey retinal visual information to the brain (Bridge et al., 2010). One putative pathway for these ‘blindsight’ abilities is the retino-tectal pathway (direct retinal projections to the superior colliculus). Another possibility is that the LGd projects to ‘secondary’ extrastriate cortical visual areas although the existence of extrastriate LGd projections has been controversial (Rabbo, Koch, Lefèvre, & Seizeur, 2015).

Conflicting reports of extrastriate LGd connections using different techniques across a variety of animal species can be found throughout the literature (**Table 1**). Karl Lashley’s early retrograde degeneration studies in the rat established the historically predominant view that the VISp is the only recipient of LGd input (K. Lashley, 1934; K. S. Lashley, 1941). In 1965, extrastriate LGd connections were first reported in the opossum (I. T. Diamond & Utley, 1963) (contrary to an earlier study (Bodian, 1935)) and later in the rabbit (Rose & Malis, 1965), but both studies seemed to gain little notice from the research field. Instead, two studies in cat would establish that the cat (but not primate) LGd provided extensive input to VISp as well as multiple extrastriate visual areas (Glickstein, King, Miller, & Berkley, 1967; Wilson & Cragg, 1967). Initially, the cat extrastriate LGd projections were thought to be an exception compared to other mammals, but both positive (L. Benevento & Yoshida, 1981; Bullier & Kennedy, 1983; Coleman & Clerici, 1980; Dräger, 1974; Fries, 1981; Garey & Powell, 1971; Haight, Sanderson, Neylon, & Patten, 1980; Hall & Diamond, 1968; Hernández-González, Cavada, & Reinoso-Suarez, 1994; Holländer & Hälbig, 1980; Hubel, 1975; Hughes, 1977; Karamanlidis, Saigal, Giolli, Mangana, & Michaloudi, 1979; Kennedy & Bullier, 1985; LeVay & Gilbert, 1976; Lysakowski, Standage, & Benevento, 1988; Niimi & Sprague, 1970; Raczkowski & Rosenquist, 1983; Ribak & Peters, 1975; Sanderson, Dreher, & Gayer, 1991; Tanaka, Lausmann, & Creutzfeldt, 1990; Towns, Burton, Kimberly, & Fetterman, 1982; Weber, Casagrande, & Harting, 1977; Winfield, Gatter, & Powell, 1975; Wong-Riley, 1976; Yukie & Iwai, 1981) and negative/absent (L. Benevento & Ebner, 1971; L. A. Benevento & Standage, 1982; Caviness & Frost, 1980; Coleman & Clerici, 1981; Coleman, Diamond, & Winer, 1977; Colwell, 1975; I. Diamond, Snyder, Killackey, Jane, & Hall, 1970; Dräger, 1981; Dürsteler, Blakemore, & Garey, 1979; Garey & Powell, 1971; Glendenning, Kofron, & Diamond, 1976; Gould, Hall, & Ebner, 1978; Harting, Diamond, & Hall, 1973; Hubel & Wiesel, 1972; Kaas, Hall, & Diamond, 1972; Karamanlidis & Giolli, 1977; Peters & Saldanha, 1976; Robson & Hall, 1975) reports of LGd extrastriate projections in multiple species led to debate over the next several decades.

**Table 1.** List of articles reporting extrastriate projections of the lateral geniculate thalamus across different species. Articles which examined thalamocortical connectivity of the LGd and/or VIS cortices are listed in chronological order. Columns include brief information about the experimental species, methodology, whether or not the study reported the existence or non-existence of LGd extrastriate connections, and a brief summary description of the reported findings. Overall, we found 30 reports of positive findings of extrastriate LGd projections, 16 reports of negative findings, and 4 studies which did not report their presence or absence (DNR). Different studies have used different nomenclature to describe visual cortices, but secondary visual cortex is located immediately adjacent and surrounding primary visual cortex in mammalian species. Generally, primary visual cortex = VISp, V1, area 17, area striata, striate cortex; secondary visual cortices = V2-V4 and MT, areas 18 (18a and 18b) and 19, area occipitalis, peristriate, extrastriate (mouse = VISal, VISl, VISpl, VISam, VISpm). AAV, adeno-associated virus; AD, anterograde degeneration; AT, anterograde tracing; LGd, dorsal lateral geniculate thalamic nucleus; MT, middle temporal visual area; RD, retrograde degeneration; RT, retrograde tracing.

| Article | Animal Model | Method | Extrastriate LGd Connections? | Summary Findings |
| --- | --- | --- | --- | --- |
| Lashley, 1934 | Rat | RD after lesions in cortex | No | No degeneration is evident except after lesions of the area striata |
| Bodian, 1935 | Opposum | RD after cortical lesions | No | Lgd degeneration after striate cortex lesions, but no other cortical area |
| Diamond and Utley, 1963 | Opposum | RD after striate/peristriate lesions | Yes | Degeneration in LGd was greater when lesions included both peristriate and striate |
| Rose and Malis, 1965 | Rabbit | RD after striate and peristriate lesions | Yes | Degeneration in ventromedial LGd following medial peristriate lesion |
| Glickstein et al, 1967 | Cat | AD after LGd lesions | Yes | Dense LGd projections to Areas 17, 18, and 19 |
| Wilson and Cragg, 1967 | Monkey and Cat | AD after thalamic lesions | Yes (cat)<br>No (monkey) | In monkey, degeneration did not extend beyond the borders of area 17 after LGd lesions. In cat, degeneration was found in area 17 and area 18 after LGd lesions |
| Hall and Diamond, 1968 | Hedgehog | RD after striate and peristriate lesions | Yes | Degeneration in LGd following extrastriate lesions |
| Diamond et al, 1970 | Tree Shrew | RD after striate and peristriate lesions | No | No degeneration in LGd following peristriate lesions |
| Niimi and Sprague, 1970 | Cat | RD after cortical lesions | Yes | Lesions of area 18 and 19 produced marked retrograde degeneration in medial interlaminar nucleus and mild degeneration in other LGd lamina |
| Benevento and Ebner, 1971 | Opposum | AD after thalamus lesions | No | Lgd lesions did not produce degeneration anywhere other than striate cortex |
| Garey and Powell, 1971 | Cat and monkey | AD after thalamic lesions | Yes (cat)<br>No (monkey) | Degeneration was found in areas 17, 18, 19, and the lateral suprasylvian area after lesions of the cat LGd. In contrast, lesions of monkey LGd produced degeneration that was restricted to area 17 |
| Hubel and Wiesel, 1972 | Macaque monkey | AD after thalamic lesions | No | No degeneration was seen outside of the striate cortex |
| Kaas et al, 1972 | Squirrel | RD after cortical lesions | No | No degeneration in LGd following Area 18 or Area 19 lesions |
| Harting et al, 1973 | Tree Shrew | AD after thalamic lesions | No | No degeneration observed outside of Area 17 after LGd lesions |
| Dräger, 1974 | Mouse | Trasneuronal AT w/ tritiated Proline into eye | Yes | Radioactivity in Area 18a, unclear if from LGd or VISp |
| Colewell, 1975 | Rabbit | RT and AT via coinjection of HRP and tritiated amino acids | DNR | Did not report any extrageniculate LGd projections |
| Hubel, 1975 | Tree Shrew | Trasneuronal AT w/ tritiated Proline into eye | Yes | Radioactivity in Area 18a, unclear if from LGd or VISp |
| Ribak and Peters, 1975 | Rat | AT w/ tritiated Proline into LGd | Yes | Radioactivity in Areas 18, and 18a, unclear if from LGd or VISp |
| Robson and Hall, 1975 | Squirrel | AT w/ tritiated amino acids into thalamus | DNR | Did not report any extrageniculate LGd projections |
| Winfield et al, 1975 | Rhesus monkey | RT w/ HRP into cortex | Yes | Description of results is vague, but in LGd of all brains there has been a well defined band of labelled cells throughout all laminae in the caudal third of its anteroposterior extent." Figure show an injection site in Area 18 that produces labeling across all layers at the medial boundary of LGd |
| Glendenning et al, 1976 | Galago bush baby | AD after thalamic lesions | No | No mention of AD anywhere other than area 17 |
| Levay and Gilbert, 1976 | Cat | AT w/ Tritiated Amino Acids into different LGd lamina | Yes | Radioactivity in area 17 produced from injections in the LGd 'A' lamina and radioactivity in areas 17, 18, 19, and the suprasylvian gyrus after injections in the LGd 'C' lamina. Distribution of projection fibers in area 18 lamina similar to area 17 organization |
| Peters and Saldanha, 1976 | Rat | AD after thalamic lesions and EM reconstruction of synaptic contacts in cortex | DNR | Did not report any extrageniculate LGd projections |
| Wong-Riley, 1976 | Squirrel monkey | RT w/ HRP after injections into dorsolateral perstriate along the area 18 and area 19 border | yes | HRP-labelled LGd cells occupied a central wedge or sector exclusively in the anterior one-third of the nucleus. This wedge extended rostralateral to caudomedially through all of the parvocellular and magnocellular laminae (see Fig. 3) and corresponded to the peripheral representation of the horizontal meridian and the immediately adjacent fields 11. Labelled almost all of the medium and large cells and none of the small neurons |
| Coleman et al, 1977 | Opposum | RT w/ HRP after injections into cortex | No | No retrograde labeled LGd neurons after HRP injections into extrastriate cortex |
| Hughes, 1977 | Rat | AT w/ tritiated amino acids after thalamus injections and RT w/ HRP after cortex injections | Yes | Tritiated amino acid injection into LGd produces radioactivity in the medial aspect of Area 18a and HRP injection into Area 18a results in extensive retrograde labelling in LGd |
| Karamanlidis and Giolli, 1977 | Rabbit | Horseradish-Peroxidase (HRP) injections into V1 cortex | DNR | Did not report any extrageniculate LGd projections |
| Weber et al, 1977 | Grey-Squirrel | Trasneuronal AT w/ tritiated Proline into eye | Yes | Radioactivity in Area 18a, unclear if from LGd or VISp |
| Gould et al, 1978 | Hedgehog | AD after thalamic lesions | No | No degeneration in parastriate cortex following LGd lesions |
| Dursteler et al, 1979 | Hamster | RT w/ HRP injected into areas 17, 18a, 18b | No | Found HRP-labeled LGd cells after injection into area 18a, but attributed labeling to leakage into area 17 |
| Karamanlidis et al, 1979 | Sheep | RT w/ HRP afterinjections into cortex | Yes | Retrogradely-labeled HRP neurons in LGd after injections in either area striata and area occipitalis |
| Coleman and Clerici, 1980 | Rat | RT w/ HRP afterinjections into cortex | Yes | Scattered, sparse HRP-labeled LGd neurons following injections in Area 18a |
| Haight et al, 1980 | Marsupial brush-tailed possum | AT w/ tritiated amino acids and RT w/ HRP after injections into cortex | Yes | Overlapping distribution of tritiated amino acid fibers and retrogradely-labeled HRP neurons within LGd. LGd projections to peristriate similar to genicul-striate projections, preferential to outer portions of LGd. |
| Hollander and Halbig, 1980 | Rabbit | Trasneuronal AT w/ tritiated amino acids into eye and RT w/ HRP after cortex injections | Yes | Tritiated amino acid labeling in area striata and area occipitalis following intraocular injection and retrograde HRP-labeled neurons in LGd following injection into area striata and area occipitalis |
| Caviness and Frost, 1980 | Mouse | AD after thalamic lesions | No | Degeneration confined to Area 17 after LGd lesions |
| Dräger, 1981 | Reeler mice | Trasneuronal AT w/ tritiated amino acids into eye | No | Radioactivity present in Area 18a, but interpreted as arising from tectum |
| Benevento and Yoshida, 1981 | Macaque monkey | AT w/ tritiated amino acids into LGd and RT w/ HRP into Areas 18 and 19 | Yes | Interlaminar LGd neurons project to Layer V in a large part of Area 19 and anterior Area 18 |
| Fries, 1981 | Macaque monkey | HRP into Areas 18 and 19 | Yes | After injections in area 18 and 19, HRP-labeled LGd cells were found broadly scattered across LGd lamina. Numbers of labelled neurons were low and never exceeded 12 per section. |
| Coleman and Clerici, 1981 | Opposum | RT w/ HRP after cortex injections | No | No HRP-labeled LGd neurons following injections into Area 18, 19, or anterior and posterior peristriate area |
| Yukie and Iwai, 1981 | Macaque monkey | RT w/ HRP after injections into striate, prestriate, inferotemporal and parietal cortices | Yes | HRP labeled LGd cells projecting to peristriate show unique distribution and morphology compared to striate-projecting LGd neurons |
| Benevento and Standage, 1982 | Macaque monkey | RT w/ wheat germ conjugated HRP into V2 and MT | No | No retrograde labeling in the LGd after injections into V2 and MT |
| Towns et al, 1982 | Rabbit | AT w/ tritiated amino acids into LGd and RT w/ HRP into striate and occipital cortex | Yes | Radioactivity in occipital cortex, confirmed by retrogradely-labeled HRP neurons in LGd after occipital cortex injection |
| Bullier and Kennedy, 1983 | Macaque monkey | RT w/ Fast Blue and Diamidino Yellow into V2 | Yes | Scattered retrograde labeling within interlaminar and S layers |
| Raczkowski and Rosenquist, 1983 | Cat | RT w/ HRP and AT w/ tritiated Leucine in multiple visual cortices | Yes | Lgd projects to areas 17, 18, 19, and suprasylvian areas |
| Kennedy and Bullier, 1985 | Macaque monkey | RT w/ Fast Blue and Diamidino Yellow into V1 and V2 | Yes | V1-projecting LGd neurons were densely packed within lamina whereas V2-projecting LGd neurons were rare, more scattered, and located within interlaminar and S layers |
| Lysakowski et al, 1988 | Macaque monkey | RT w/ Granular Blue and Nuclear Yellow into V1 and V4 | Yes | V4-projecting LGd neurons were sparse compared to V1-projecting LGd neurons. No collateralization of input from LGd to V1 and V4 |
| Tanaka et al, 1990 | Macaque monkey | RT w/ HRP after cortex injections | Yes | Sparse numbers of V4-projecting LGd neurons in the interlaminar region |
| Sanderson et al, 1991 | Rat | RT w/ WGA-HRP after cortex injections | Yes | Minor retrograde labeling (less than 5%) in LGd after injections in AM, AL, and LM |
| Hernandez-Gonzalez et al, 1994 | Macaque monkey | RT w/ multiple retrograde tracers into inferior temporal cortex | Yes | Sparse numbers of LGd interlaminar neurons project to inferior temporal cortex, appear similar to population of other extrastriate LGd neurons |
| Oh et al, 2014 | Mouse | AT w/ fluorescent AAV vectors | Yes | Lgd sends output to VISp, VISal, and VISl and LGd receives input from VISam, VISp, VISal, VISl, and the temporal association area |

With the advent of new tract tracing methods in the 1980’s, the existence of LGd projections to extrastriate visual cortex became more established in non-human primates. LGd axons have been reported within V2 (L. Benevento & Yoshida, 1981; Bullier & Kennedy, 1983; Fries, 1981; Kennedy & Bullier, 1985; Yukie & Iwai, 1981), V4 (Lysakowski et al., 1988; Tanaka et al., 1990), inferotemporal cortex (Hernández-González et al., 1994), and MT, although some studies reported a lack of fibers in these areas (L. A. Benevento & Standage, 1982). Non-human primate extrastriate LGd projection neurons are scattered throughout the interlaminar and S layers but are larger in size than typical koniocellular neurons (Bullier & Kennedy, 1983; Hendry & Reid, 2000). Using V1-lesioned macaque monkeys, Schmid and colleagues found that reversible inactivation of the LGd in VISp-lesioned animals eliminated extrastriate cortex fMRI responses and ‘blindsight’ behavior, proposing that LGd extrastriate connections mediate blindsight and may provide a shortcut for rapid detection in normal vision (Schmid et al., 2010).

Compared to the non-human primate, the rodent LGd has been significantly less studied. However, the popularity of the mouse as a genetic model has renewed interest in rodent vision (Ringach et al., 2016; Seabrook, Burbridge, Crair, & Huberman, 2017), particularly toward discovering parallel visual processing streams through the LGd (Denman & Contreras, 2016; Kerschensteiner & Guido, 2017; Morgan, Berger, Wetzel, & Lichtman, 2016). A major obstacle to understanding the organization of the visual system in rodents has been structural differences in the visual thalamus and cortex. Notably, the lack of a laminar structure to the rodent LGd hinders a clear understanding of LGd cell type organization. In addition, it is unclear how extrastriate visual cortex regions in the mouse are homologous to the more complex non-human primate. A recent mouse connectomics study reported multiple LGd extrastriate connections, but did not specifically address this topic (Oh et al., 2014). As part of our Mouse Connectome Project (www.mouseconnectome.org), we have performed double anterograde/retrograde tracer coinjections across the cortex, including VISp and the medially (VISam, VISpm) and laterally adjacent extrastriate VIS areas (VISal, VISl, and VISpl) (Dong, 2008). To understand VIS thalamocortical connectivity and investigate LGd extrastriate connectivity in the mouse, we have targeted anterograde/retrograde tracer coinjections into VIS cortices and small subregions of the LGd thalamus. By simultaneously visualizing input/output connectivity, this approach can address bidirectional LGd thalamocortical connectivity with two cross-validated datasets (**Fig. 1**).

**Figure 1.**
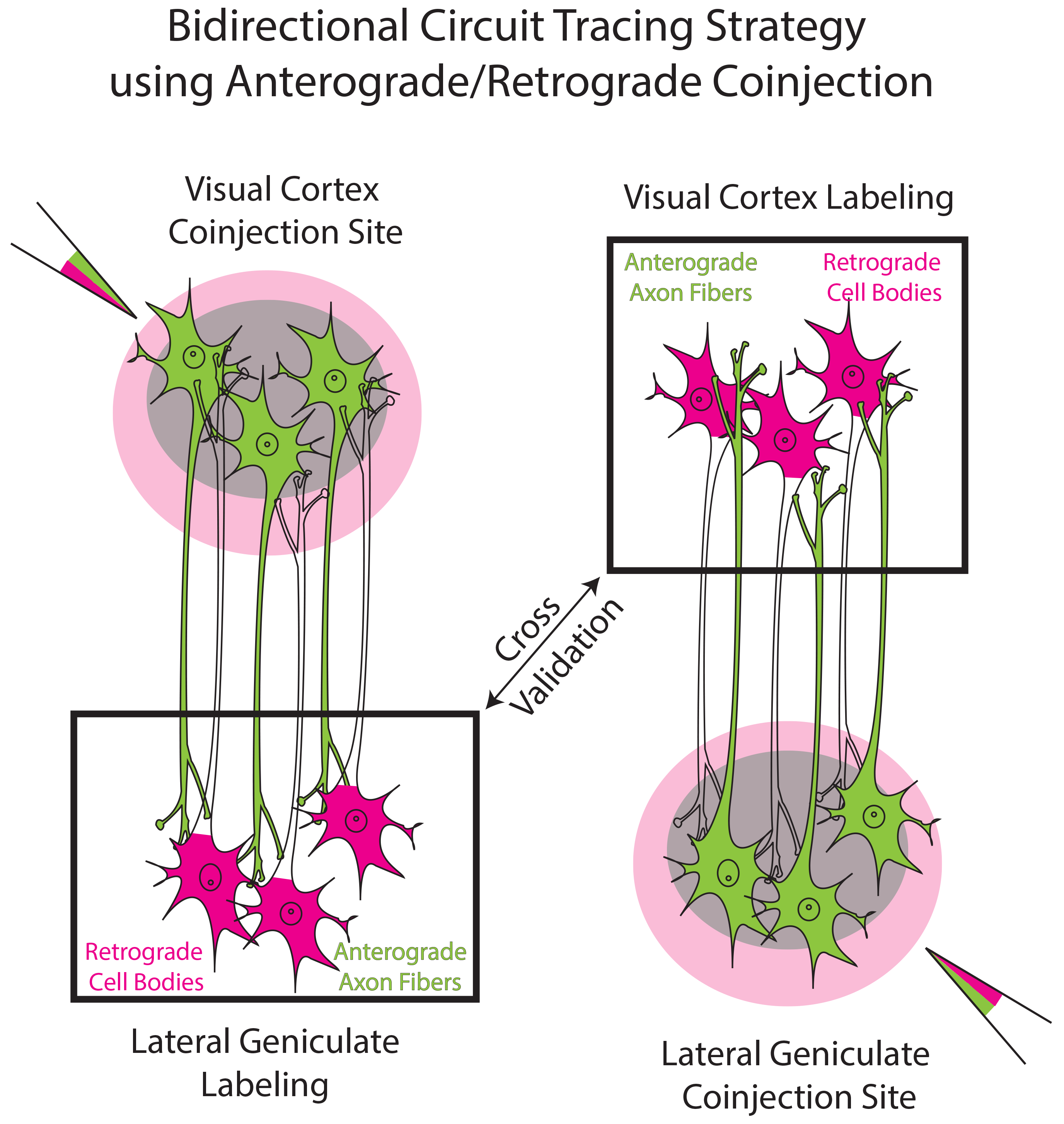
Bidirectional circuit tracing strategy. To examine bidirectional connectivity between the LGd and VIS cortices, we iontophoretically coinjected pairs of anterograde and retrograde tracers (PHAL/CTB or BDA/FG) into the LGd and visual cortices (VISp, medial extrastriate VIS areas (VISam, VISpm), and lateral extrastriate visual areas (VISal, VISl, VISpl)). (left) Coinjection sites within the VIS cortices produced anterogradely-labeled fibers (green) and retrogradely-labeled cell bodies (magenta) within the LGd. Conversely, LGd coinjections produced anterogradely-labeled fibers (green) and retrogradely-labeled cell bodies (magenta) in the VIS cortices (right). In this way, the same connection can be labeled by an anterograde tracer or a retrograde tracer and both datasets are cross-validated by the different tracer type (i.e., anterogradely-labeled LGd fibers in the visual cortices are confirmed by retrograde labeling in the LGd after VIS cortex injection). BDA, biotinynlated dextran amine; CTB, cholera toxin subunit B; FG, Fluorogold; LGd, dorsal lateral geniculate thalamic nucleus; PHAL, *phaseolus vulgaris* leucoagglutinin; VISam, anteromedial visual cortex; VISal, anterolateral visual cortex, VISl, lateral visual cortex, VISp, primary visual cortex; VISpl, posterolateral visual cortex; VISpm, posteromedial visual cortex.

## Materials and Methods

Mouse Connectome Project (MCP) tract-tracing data was generated within the Center for Integrative Connectomics (CIC) at the University of Southern California (USC) Mark and Mary Stevens Neuroimaging and Informatics Institute. MCP experimental methods and online publication have been described previously (Hintiryan et al., 2016; Zingg et al., 2014). All MCP tract-tracing experiments were performed using 8-week old male C57BL/6J mice (Jackson Laboratories). Mice had ad libitum access to food and water and were pair-housed within a temperature-(21-22°C), humidity-(51%), and light-(12hr:12hr light/dark cycle) controlled room within the Zilkha Neurogenetic Institute vivarium. All experiments were performed according to the regulatory standards set by the National Institutes of Health Guide for the Care and Use of Laboratory Animals and by the institutional guidelines described by the USC Institutional Animal Care and Use Committee.

### Tracer injection experiments

The MCP uses a variety of combinations of anterograde and retrograde tracers to simultaneously visualize multiple anatomical pathways within the same Nissl-stained mouse brain. The standard experimental approach is a double coinjection of paired anterograde (2.5% *phaseolus vulgaris* leucoagglutinin (PHAL; Vector Laboratories), 5% biotinylated dextran amine (BDA; Invitrogen) and retrograde (0.25% Alexa Fluor 647 conjugated cholera toxin subunit b (CTB-647; Invitrogen) or 1% Fluorogold (FG; Fluorochrome, LLC)) tracer into two different brain regions. Additionally, quadruple retrograde experiments are performed using 1% FG and 0.25% Alexa Fluor-488, −555, and −647 conjugated CTB tracers (CTB-488, CTB-555, CTB-647) injected into four distinct brain regions. In this study, we describe coinjection experiments in the LGd and visual cortex (VISp, VISam, VISpm, VISal, and VISl) as well as a quadruple retrograde tracer experiment in the visual cortices.

### Stereotaxic Surgeries

On the day of experiment, mice are deeply anesthetized and mounted into a Kopf stereotaxic apparatus where they are maintained under isoflurane gas anesthesia (Datex-Ohmeda vaporizer). For anterograde/retrograde coinjection experiments, tracer cocktails were iontophoretically delivered via glass micropipettes (outer tip diameter 15-30μm) using alternating 7 s on/off pulsed positive electrical current (Stoelting Co. current source) for 5 (BDA or AAV/FG) or 10 min (PHAL/CTB-647). For quadruple retrograde tracing experiments, 50nl of retrograde tracers were individually pressure-injected via glass micropipettes at a rate of 10nl/min. All injections were placed in the right hemisphere. Injection site coordinates for the each surgery case are on the Mouse Connectome Project iConnectome viewer (www.MouseConnectome.org). Following injections, incisions were sutured and mice received analgesic pain reliever and were returned to their home cages for recovery.

### Histology and Immunohistochemical Processing

After 1-2 weeks post-surgery, each mouse was deeply anesthetized with an overdose of sodium pentobarbital and trans-cardially perfused with 50ml of 0.9% saline solution followed by 50ml of 4% paraformaldehyde (PFA, pH 9.5). Following extraction, brain tissue was postfixed in 4% PFA for 24-48hr at 4°C. Fixed brains were embedded in 3% Type I-B agarose (Sigma-Aldrich) and sliced into four series of 50μm thick coronal sections using a Compresstome (VF-700, Precisionary Instruments, Greenville, NC) and stored in cryopreservant at −20°C. For double coinjection experiments, one series of tissue sections was processed for immunofluorescent tracer localization. BDA immunofluorescence was visualized using a 647-or 568-conjugated Streptavidin. For PHAL immunostaining, sections were placed in a blocking solution containing normal donkey serum (Vector Laboratories) and Triton X (VWR) for 1 hr. After rinsing in buffer, sections were incubated in PHAL primary antiserum (donkey serum, Triton X, 1:100 rabbit anti-PHAL antibody (Vector Laboratories) in KPBS buffer solution) for 48-72 hrs. at 4°C. Sections were then rinsed again in buffer solution and then immersed in secondary antibody solution (donkey serum, Triton X, and 1:500 donkey anti-rabbit IgG conjugated with Alexa Fluor 488. Finally, all sections were stained with Neurotrace Blue for 2-3 hrs. to visualize cytoarchitecture. After processing, sections were mounted onto microscope slides and coverslipped using 65% glycerol.

### Imaging and Online Data Publication

Complete tissue sections were scanned using a 10X objective lens on an Olympus VS120 slide scanning microscope. Each tracer was visualized using appropriately-matched fluorescent filters and whole tissue section images were stitched from tiled scanning into VSI image files. For online publication, raw images are corrected for correct left-right orientation and matched to the nearest Allen Reference Atlas level (ARA; (Dong, 2008)). VSI image files are converted to TIFF file format and warped and registered to fit ARA atlas levels (all images shown in this manuscript are from unwarped, unregistered VSI images). Each color channel is brightness/contrast adjusted to maximize labeling visibility (Neurotrace blue is converted to brightfield) and TIFF images are then converted to JPEG2000 file format for online publication in the Mouse Connectome Project iConnectome viewer (www.MouseConnectome.org).

## Results

Overall, all tracer experiments into the cortex produced 8 anterograde and 9 retrograde VISp injection sites, 2 anterograde and 3 retrograde VISam injection sites, 3 anterograde and 4 retrograde VISpm injection sites, 4 anterograde and 2 retrograde VISal injection sites, and 3 anterograde and 3 retrograde VISl injection sites. Double coinjection experiments in the LGd produced 9 anterograde and 10 retrograde sites. An additional 2 anterograde/retrograde coinjections were made into the LP for comparison. Microscopy images of the entire tissue series for each experimental case are available online at www.MouseConnectome.org. The spread of each coinjection site (except LP) was mapped onto ARA sections and the overall distribution of the injection sites covered a large amount of the whole VIS cortices and LGd (**Fig. 2**).

**Figure 2.**
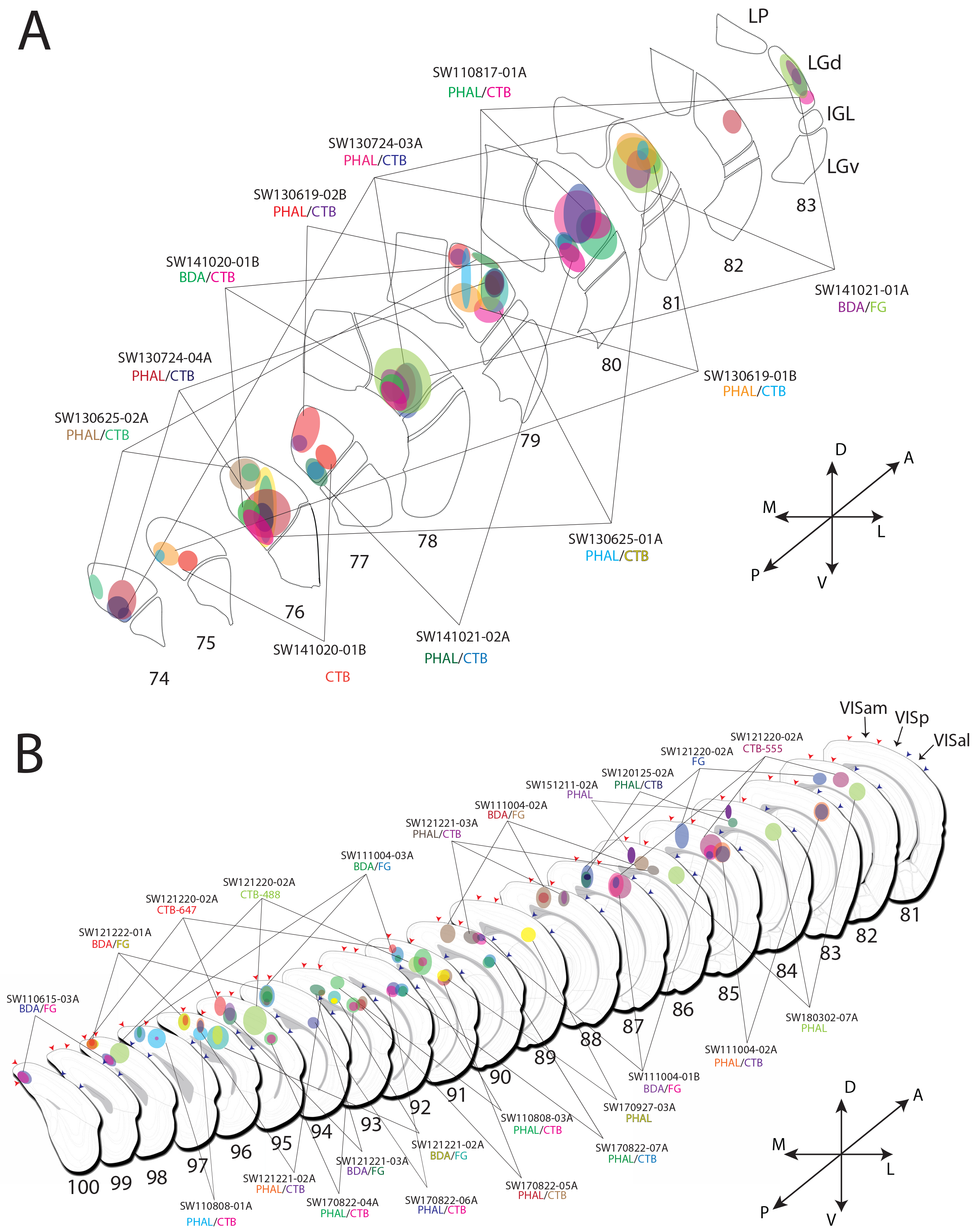
Mapped injection site spread throughout the visual cortex and thalamus. For cortical **(A)** and LGd injections **(B)**, the size and shape of each tracer injection site was mapped onto ARA sections using one 200μm-interval tissue series (ARA atlas numbers listed below each section). Borders between the VISp and medial (VISam/VISpm) and lateral extrastriate areas (VISal, VISl, VISpl) are demarcated by red and blue arrowheads, respectively. The boundaries of the LGd and neighboring LP, IGL, and LGv are labeled at the far right and apply to each consecutive rostrocaudal level. In most cortical injection cases, retrograde injection sites are spherical in shape with a 400-500μm diameter spreading across 2-3 rostrocaudal levels. Anterograde injection spread was typically smaller than retrograde spread when coinjected together. In LGd-injected experiments, coinjection sites are spherical in shape with a 200-300μm diameter spreading across 2-3 rostrocaudal levels and small enough to not completely fill the LGd (most of the injection site is concentrated at the center with minimal spread across levels).Tracer name is listed under each case number and color-coded to injection site mapped color. ARA, Allen Reference Atlas; BDA, biotinynlated dextran amine; CTB-647, cholera toxin subunit B conjugated to Alexa Fluor 647; FG, Fluorogold; IGL, intergeniculate leaflet; LGd, dorsal lateral geniculate thalamic nucleus; LP, lateral posterior thalamic nucleus; PHAL, *phaseolus vulgaris* leucoagglutinin; VISam, anteromedial visual cortex; VISal, anterolateral visual cortex, VISl, lateral visual cortex, VISp, primary visual cortex; VISpl, posterolateral visual cortex; VISpm, posteromedial visual cortex.

### Verification of VIS cortex boundaries by gene expression, cytoarchitecture, and other connectivity patterns

A critical factor in interpreting anatomical tract tracing data is the verification of injection site and tracer labeling neuroanatomical locations. Directly registering experimental tissue sections to standardized brain atlases (such as the ARA) can be difficult due to variability in histological processing across multiple animals (i.e. oblique tissue sectioning angles, tissue warping, etc.). To confirm the ARA boundaries of the VISp with the medial and lateral extrastriate regions, we examined gene expression, cytoarchitecture, and other connectivity data that would identify the VISam/VISpm and VISal/VISl/VISpl from the VISp. First, VISam and VISpm have a thinner layer 4 compared to the laterally-adjacent VISp and the medially-adjacent agranular retrosplenial cortex (RSP) that does not contain layer 4. *Rorb* gene expression has been used as a genetic marker for layer 4 and is densely expressed in layer 4 neurons across the cortex with more sparse expression in layer 5 (Harris et al., 2014). *Rorb* gene expression in the Allen Brain Atlas *in situ* hybridization image database provides a clear demarcation of the VISam and VISpm boundaries that is in strong agreement with the ARA boundary delineation (**Fig. 3**).

**Figure 3.**
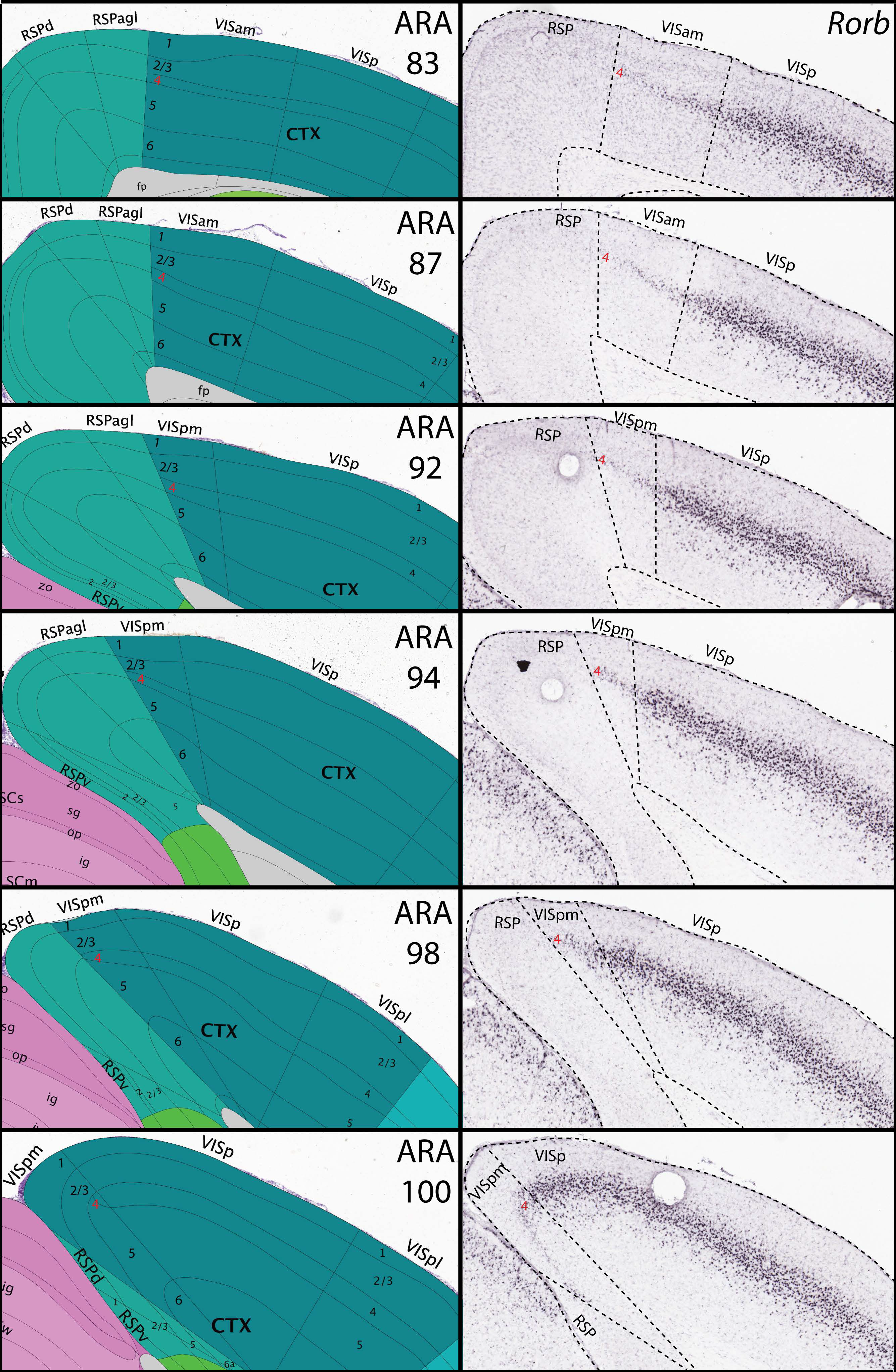
Validation of the VISam and VISpm boundaries by *Rorb* gene expression. The VISam and VISpm can be distinguished from the medially-adjacent RSP and laterally-adjacent VISp by distinct changes in layer 4. Layer 4 is absent from RSP and thicker in VISp compared to VISam and VISpm. To clearly visualize layer 4, we compared our boundaries to a layer 4 gene expression marker *(Rorb)* in the Allen Brain Atlas *in situ* hybridization database (http://mouse.brain-map.org/, ARA atlas sections on left, corresponding *Rorb in situ* hybridization on right). As expected, *Rorb* expression is robust in VISp layer 4 with more sparse expression in layer 5. In the ARA-defined VISam and VISpm, *Rorb* expression is notably weaker, in a thinner lamina compared to VISp, and abruptly ends at the RSP border across all rostrocaudal levels. ARA, Allen reference atlas; *Rorb*, RAR-related orphan receptor beta; RSP, retrosplenial cortex; RSPagl, agranular retrosplenial cortex; RSPd, dorsal retrosplenial cortex; RSPv, ventral retrosplenial cortex; VISam, anteromedial visual cortex; VISp, primary visual cortex; VISpl, posterolateral visual cortex; VISpm, posteromedial visual cortex.

For further verification of extrastriate boundaries, we analyzed extrastriate connectivity with VISp and thalamus. In the mouse, a defining characteristic of extrastriate visual areas are topographic connection with VISp (Wang & Burkhalter, 2007). In case SW121221-02A, we performed a double coinjection of BDA/FG into the lateral VISp and PHAL/CTB into the medial VISp (**Fig. 4A**). Both coinjections produced topographically-organized columns of anterograde and retrograde labeling within the ARA-defined VISam and VISal. Furthermore, these connectivity patterns are consistent with the definition of extrastriate regions as demonstrated by Wang and Burkhalter in which extrastriate regions receive topographic maps of VISp input (although it is currently unclear how to directly relate the entire coronal ARA delineation to the flattened laminar surface sectioning used by Wang and Burkhalter (Wang & Burkhalter, 2007)). In addition, ‘higher order’ thalamic nuclei such as the LP project more densely to the extrastriate visual areas than the VISp and show different laminar projection patterns (Zhou, Masterson, Damron, Guido, & Bickford, 2018). In case SW141021-02A, we coinjected BDA/FG into the caudal medial LP while PHAL/CTB was coinjected into the LGd (**Fig. 4B**). As expected, the anterograde and retrograde labeling patterns from the LP and LGd injection sites clearly distinguish the extrastriate visual areas. LP coinjection produced robust axon terminal fields in layer 4 of the ipsilateral extrastriate visual areas and more minor fiber labeling in VISp layer 5a. Retrogradely-labeled LP-projecting neurons were densely distributed in layers 5, 6a, and 6b in extrastriate areas whereas retrograde labeling in VISp was limited to layers 5 and 6b.

**Figure 4.**
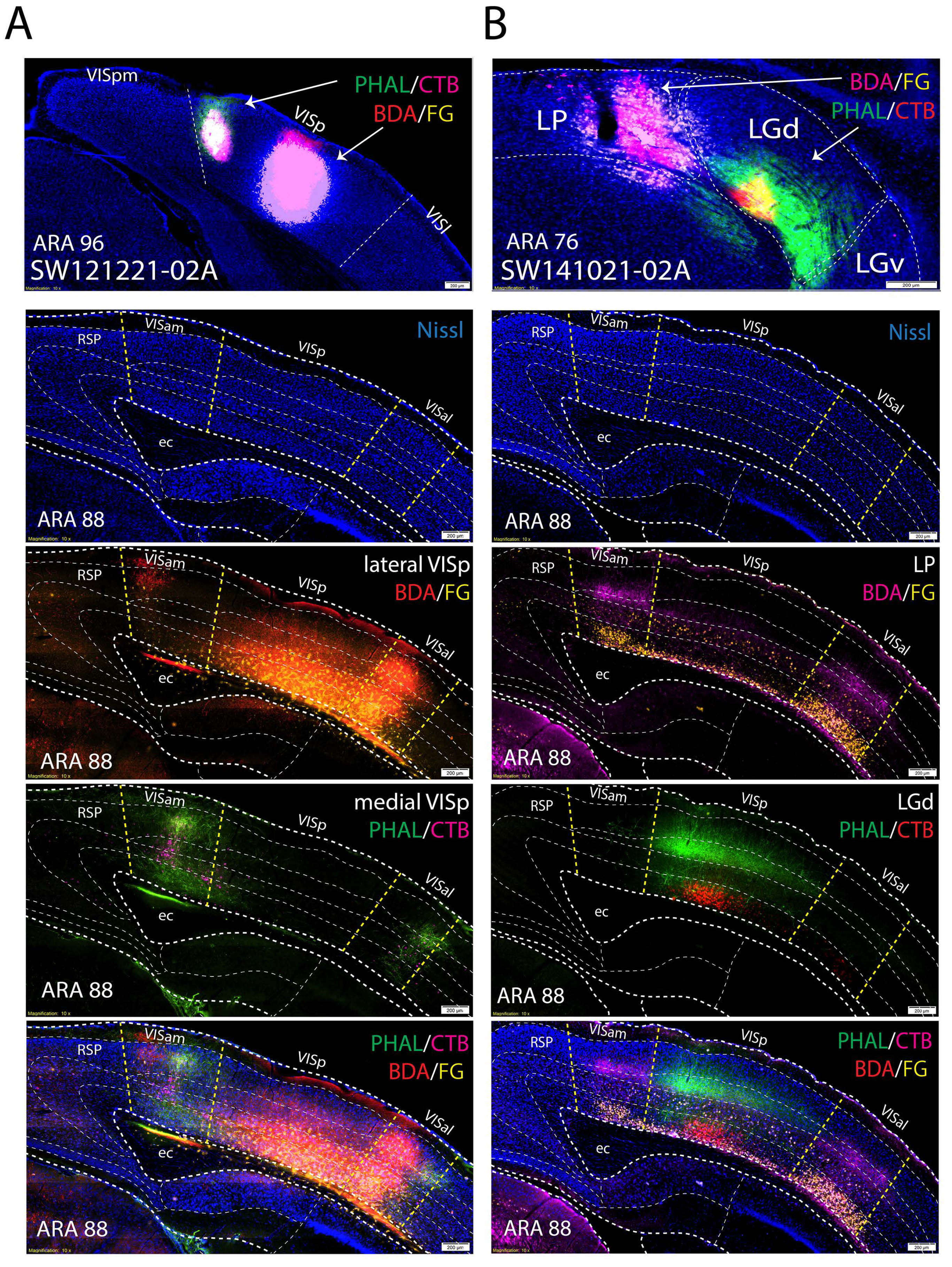
Validation of the VISam and VISpm boundaries by *Rorb* gene expression. **(A)** VISp is distinguished from extrastriate visual areas by differences in anatomical connectivity. A double coinjection into the medial and lateral parts of VISp (SW121221-02A) reveals the bidirectional topographic connections in the VISam and VISal. BDA/FG coinjection into lateral VISp produced a column of anterogradely-labeled fibers (red) and retrogradely-labeled neurons (yellow) in the medial parts of VISam and VISal. PHAL/CTB coinjection into the medial VISp produced a column of anterogradely-labeled fibers (green) and retrogradely-labeled neurons (magenta) in the lateral parts of VISam and VISal. Together, the full set of topographic connections are within the VISam and VISal boundaries as demarcated in the ARA (bottom). **(B)** Double coinjection into the thalamus (SW141021-02A) also reveals bidirectional connectivity features that are unique to extrastriate regions compared to VISp and are consistent with ARA boundaries. BDA/FG coinjection into caudal medial part of LP produced thick anterogradely-labeled fiber terminal fields (magenta) in layer 4 of VISam and VISal whereas only superficial layer 5 was innervated in VISp. In addition, retrogradely-labeled neurons were located in layers 5, 6a, and 6b in VISam and VISal whereas retrograde labeling was absent from VISp layer 6a. Coinjection of PHAL/CTB into the LGd produced anterogradely-labeled fiber terminal fields (green) in VISp layer 4 and retrogradely-labeled neurons (red) in layer 6a that were mostly localized to the VISp region. Together, the set of connections show clear differences in connectivity that distinguish the extrastriate visual areas from the VISp. ARA, Allen Reference Atlas; BDA, biotinynlated dextran amine; CTB, cholera toxin subunit B; ec, external capsule; FG, Fluorogold; IGL, intergeniculate leaflet; LGd, dorsal lateral geniculate thalamic nucleus; LP, lateral posterior thalamic nucleus; PHAL, *phaseolus vulgaris* leucoagglutinin; RSP, retrosplenial cortex; VISam, anteromedial visual cortex; VISal, anterolateral visual cortex, VISl, lateral visual cortex, VISp, primary visual cortex; VISpl, posterolateral visual cortex; VISpm, posteromedial visual cortex.

### Distribution of tracer labeling within LGd after VISp vs. VISam/VISpm coinjections

Each VISp coinjection produced a highly overlapped distribution of ipsilaterally clustered anterograde and retrograde labeling within the LGd (**Fig. 5**). Tracer labeled clusters extended across multiple rostrocaudal levels of the LGd, reflecting the spherical spread of the VIS cortex injection site. As expected, the overall rostrocaudal and mediolateral locations of the VISp injection sites demonstrated that the clusters of labeling within the LGd were topographically-organized. As shown by the labeling distribution in **Fig. 5**, injection sites which were located in the posterior VISp produced retrograde labeling along the superficial, lateral edge of the LGd. In contrast, more anterior VISp injections sites retrogradely labeled LGd neurons closer to the deep, medial LGd border. In the mediolateral direction, lateral VISp injection sites produced labeling clusters close to the dorsal LGd border (adjacent to the lateral posterior thalamic nucleus (LP)), whereas more medial injection sites retrogradely-labeled LGd neurons more ventrally. However, coinjections within the most medial parts of VISp did not produce labeling at the most ventral part of the LGd (adjacent to the intergeniculate leaflet, IGL) as would be expected based on the principles of topographic organization.

**Figure 5.**
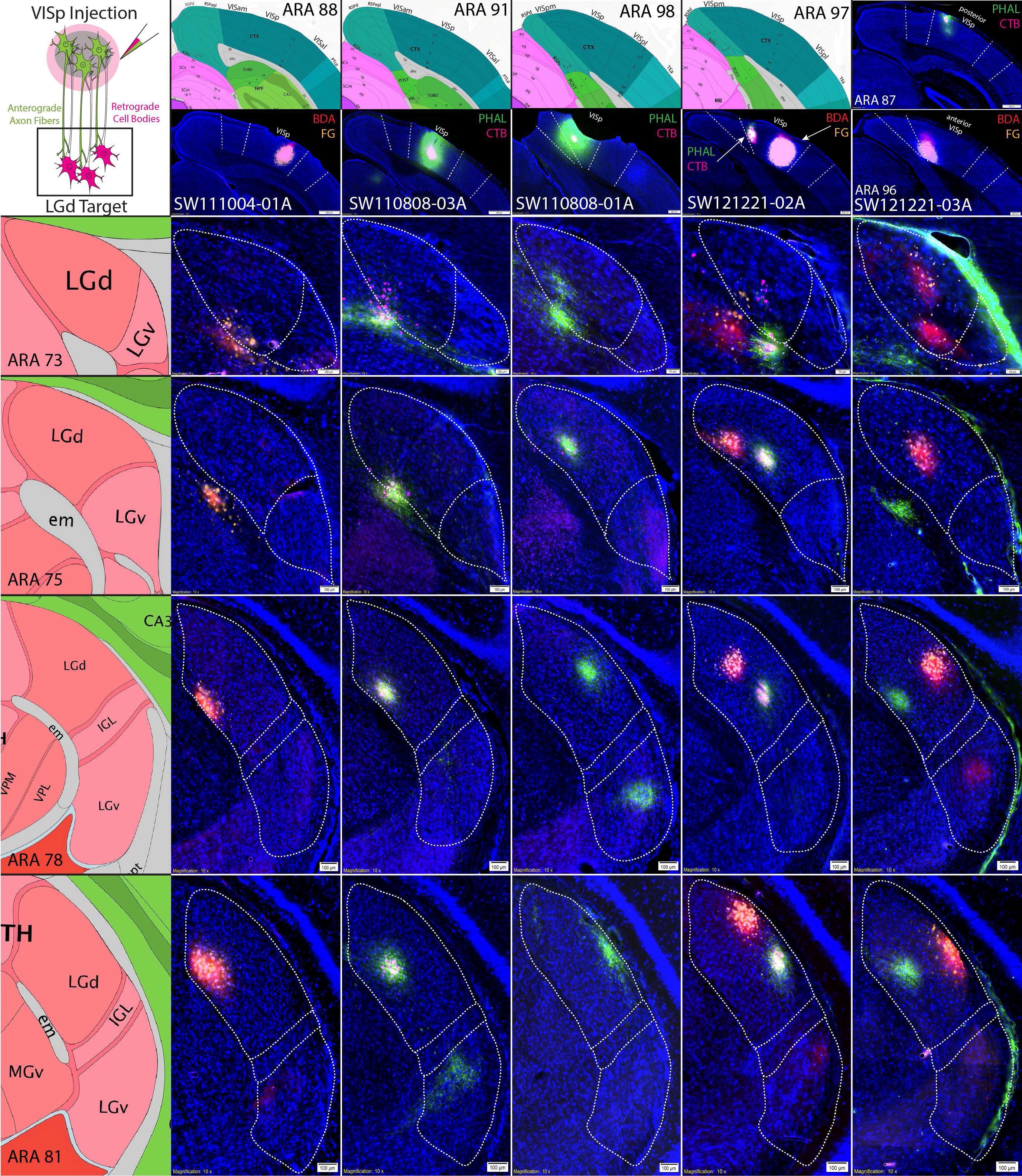
VISp coinjections show specific topographic bidirectional connectivity within LGd. Five anterograde/retrograde tracer coinjections into different parts of the VISp are shown (columns). Coinjection site center is shown at the top of each column below the corresponding ARA level. For each injection case, there are dense clusters of overlapping anterogradely-labeled fibers/retrogradely-labeled cell bodies at each rostrocaudal level of the LGd (rows). Coinjection sites that are located more caudal within the VISp produced tracer labeling clusters that are distributed along the superficial LGd (lateral) whereas bidirectional labeling clusters from more rostral VISp coinjections is in deeper parts of the LGd (medial). Coinjection sites that are located in more lateral VISp produced labeling clusters near the dorsomedial LGd/LP border whereas more medial VISp coinjections are distributed more ventromedial parts of LGd. ARA, Allen Reference Atlas; BDA, biotinynlated dextran amine; CTB, cholera toxin subunit B; FG, Fluorogold; IGL, intergeniculate leaflet; LGd, dorsal lateral geniculate thalamic nucleus; LGv, ventral lateral geniculate thalamic nucleus; LP, lateral posterior thalamic nucleus; PHAL, *phaseolus vulgaris* leucoagglutinin; VISam, anteromedial visual cortex; VISal, anterolateral visual cortex, VISl, lateral visual cortex, VISp, primary visual cortex; VISpm, posteromedial visual cortex.

Instead, coinjections into the VISam and VISpm resulted in distributed tracer labeling within the LGd along a ventral strip adjacent to the IGL border (**Fig. 6**). Similar to tracer labeling following VISp injections, coinjections within the VISam and VISpm produced small localized clusters of anterograde and retrograde labeling within the LGd. Mapping the distribution of retrogradely-labeled LGd neurons produced by the array of injection sites along the rostrocaudal extent of the VISam and VISpm determined that posterior VISpm injections retrogradely-labeled neurons at the most superior, ventrolateral part of the LGd. Coinjection sites which were located progressively more rostral through the VISam produced labeled neuron clusters more ventromedially along the LGd border adjacent to IGL.

**Figure 6.**
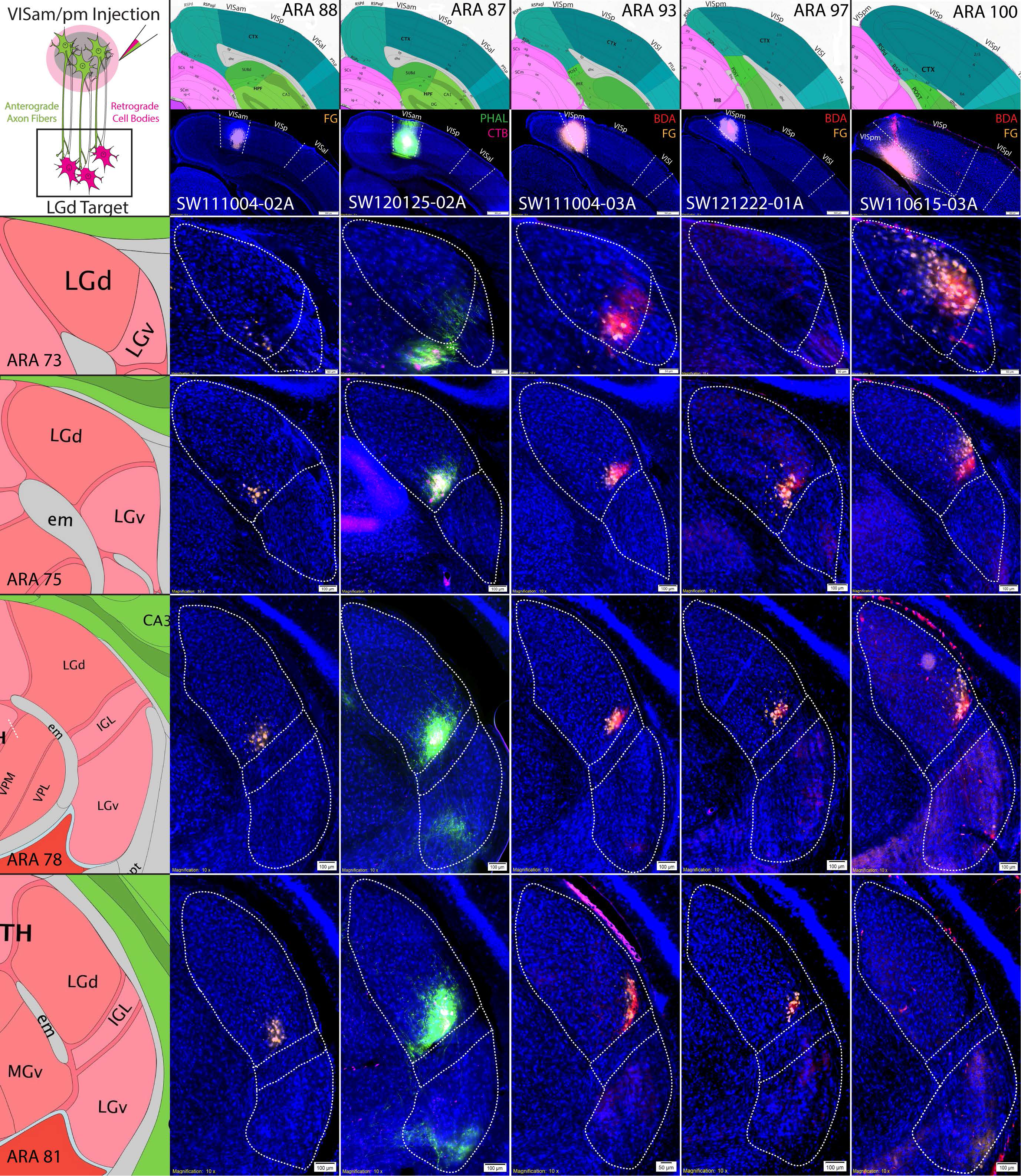
VISam and VISpm coinjections produce labeling clusters along the ventromedial LGd border. Six anterograde/retrograde tracer coinjections into VISam or VISpm are shown (columns). Coinjection site center is shown at the top of each column below the corresponding ARA level. Similar to VISp coinjections, VISam and VISpm coinjections produce dense clusters of overlapping anterogradely-labeled fibers/retrogradely-labeled cell bodies that are located in a small strip along the ventromedial LGd border adjacent to the IGL at each rostrocaudal level of the LGd (rows). Labeled clusters from VISam coinjections are distributed closer to the ventromedial corner of the LGd whereas labeled clusters from VISpm coinjections are distributed closer to the ventrolateral corner of the LGd. ARA, Allen Reference Atlas; BDA, biotinynlated dextran amine; CTB, cholera toxin subunit B; FG, Fluorogold; IGL, intergeniculate leaflet; LGd, dorsal lateral geniculate thalamic nucleus; LGv, ventral lateral geniculate thalamic nucleus; LP, lateral posterior thalamic nucleus; PHAL, *phaseolus vulgaris* leucoagglutinin; VISam, anteromedial visual cortex; VISal, anterolateral visual cortex, VISl, lateral visual cortex, VISp, primary visual cortex; VISpm, posteromedial visual cortex.

To compare the distribution of VISp and VISam/pm labeling within the same animal, we used a quadruple retrograde tracing approach and individually injected four retrograde tracers into the anterior VISp, posterior VISp, VISam, and VISpm (**Fig. 7A**). As expected, four distinct clusters of retrograde labeling were observed within the LGd in a rectangular orientation consistent with the injection site placement. Notably, the rectangular orientation of the retrograde labeling changes at different LGd rostrocaudal levels. At mid-rostrocaudal LGd levels, the LGd is split relatively evenly by the midline between anterior VISp/VISam and posterior VISp/VISpm cortex labeling. However, this anterior/posterior midline shifts laterally in anterior LGd levels and medially in posterior LGd levels, suggesting that posterior VIS cortices are more represented at anterior LGd levels and vice versa. Across all of our experiments, mapping retrograde labeling patterns produced by VISp vs. VISam/pm coinjection sites across all LGd levels shows two distinct distributions within the LGd (**Fig. 7B**). Overall, the distribution of labeling following VISam and VISpm coinjections suggests that LGd thalamocortical topography extends across the VISp border and into the VISam/pm.

**Figure 7.**
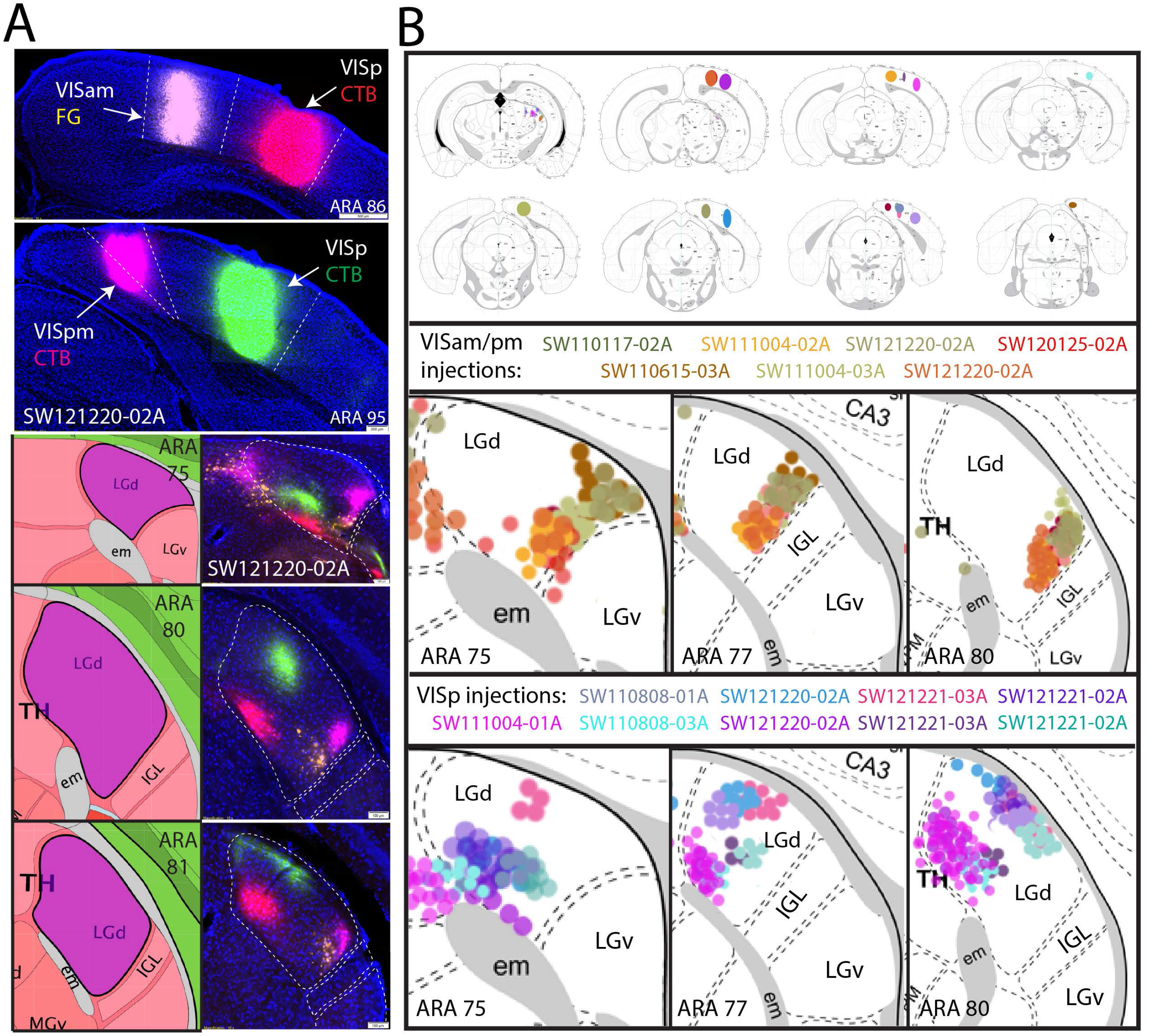
Organization of VISp vs. VISam/pm thalamocortical connectivity. **(A)** Quadruple retrograde tracer experiment to simultaneously view LGd thalamocortical connectivity with VISp, VISam, and VISpm. (top) A FG injection into the VISam (yellow) with CTB-555 injection into the anterior VISp (red) at the same rostrocaudal level. Within the same mouse, CTB-647 injection into the VISpm (magenta) and CTB-488 injection (green) into the posterior VISp at the same rostrocaudal level. (below) The distribution of the four retrograde tracers within the LGd alongside representative ARA levels are shown. Within the rostral LGd, labeling from the posterior VIS cortices has a greater representation whereas anterior VIS cortices are more represented at caudal LGd levels. At the middle LGd level, the anterior/posterior axis is almost perfectly symmetrical within the LGd. Note, the relative spacing of all 4 tracer injection sites in the cortex is relatively maintained within the LGd. **(B)** Mapping of retrograde labeling across all injection cases reveals that VISam/pm labeling are distributed uniquely along the ventral strip of the LGd whereas VISp-projecting neurons are absent from the ventromedial border (except a small area of the ventromedial LGd corner at middle rostrocaudal levels)but are located throughout the rest of the LGd. Retrograde injection site centers were mapped onto ARA sections and the distribution and relative density of retrograde labeling in the LGd was mapped with Adobe Photoshop. ARA, Allen Reference Atlas; CTB, cholera toxin subunit B; FG, Fluorogold; IGL, intergeniculate leaflet; LGd, dorsal lateral geniculate thalamic nucleus; LGv, ventral lateral geniculate thalamic nucleus; LP, lateral posterior thalamic nucleus; PHAL, *phaseolus vulgaris* leucoagglutinin; VISam, anteromedial visual cortex; VISp, primary visual cortex; VISpm, posteromedial visual cortex.

### Distribution of tracer labeling within VIS cortical areas after LGd coinjections

While LGd tracer labeling from cortical injection sites appears to demonstrate extrastriate-projecting LGd neurons, it’s possible that LGd labeling after VISam/VISpm injection sites was caused by spread into the adjacent medial VISp. To cross validate our findings based on cortical injection sites, we placed small iontophoretic coinjections within distinct LGd subregions (**Figs. 8, 9**). Although the ventral strip region of LGd is too small of a target to place an entirely restricted tracer deposit, coinjection sites that overlap the ventral strip region should produce tracer labeling within the VISp that spreads beyond the medial border into the VISam/VISpm. In contrast, LGd coinjection sites that do not spread into the ventral strip region should produce VISp tracer labeling patterns that do not cross the border into VISam/VISpm and are distributed according to topographic organization. Indeed, each LGd coinjection site produced labeling within distinct parts of VISp, consistent with a discrete placement of tracer deposit in a small area rather than the whole LGd (**Figs. 8, 9**). All LGd coinjections produced dense anterograde labeling within cortical layer 4 with lesser labeling extending into layers 1 and 6 and robust retrograde labeling within cortical layer 6a. For example, SW130619-01 contains a coinjection site that is located in the central part of LGd and produces a column of PHAL and CTB labeling that is similarly central along the rostrocaudal VISp (**Fig. 8**). Comparatively, case SW141021-01A and SW141021-02A contain coinjection sites within LGd closer to the IGL border but do not spread into the ventral strip region. In the cortex, tracer labeling is distributed more medially within the VISp and extend completely up to the VISam/VISpm border. In contrast, LGd coinjection sites that were located within the ventral strip region produced tracer labeling that was distributed in the medial VISp and also in the VISam and VISpm at multiple rostrocaudal levels (**Fig. 9**). For comparison, SW140827-01A contains a coinjection within the caudal lateral LP that produces labeling that is distributed along the VISam/VISpm rostrocaudal axis and outlines the location of the VISam/VISpm.

**Figure 8.**
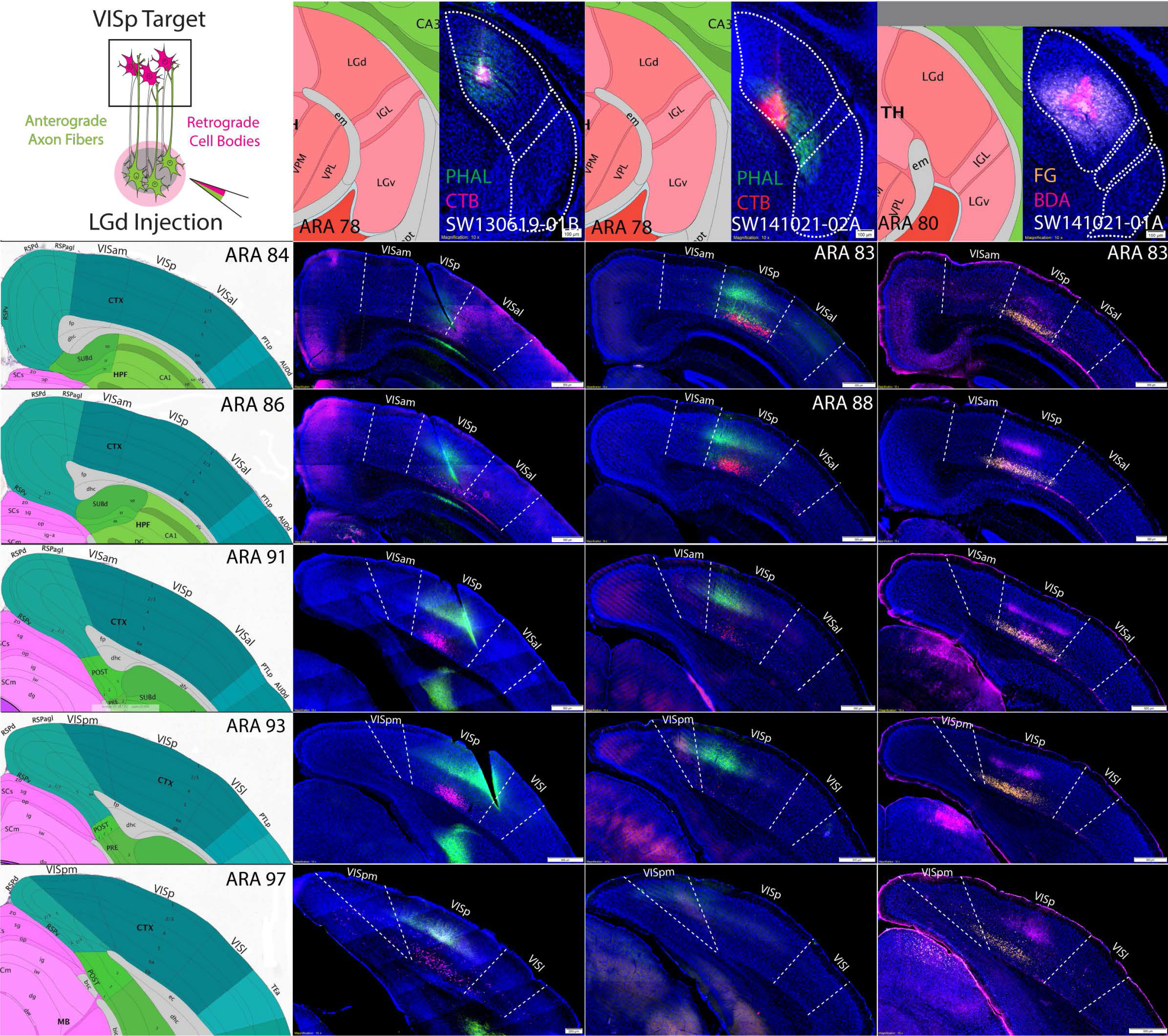
Anterograde and retrograde labeling pattern within VISp after coinjections into the LGd that
avoid the ventral strip region. Three coinjections of anterograde/retrograde tracer into different subregions of the LGd (columns) that avoid the ventral strip region produce anterogradely-labeled fibers and retrograde labeling in the VISp, but not the VISam/pm (rows). For each column, LGd tracer coinjection site is shown at the top with adjacent ARA atlas level. For each row across, the distribution of labeling throughout the rostrocaudal VIS cortices is shown (representative ARA level shown at left). In all cases, retrogradely-labeled neurons are distributed within layer 6 while anterogradely-labeled fibers are distributed in layers 1, 4 and 6 with the greatest density in layer 4. In case SW130619-01B, the LGd coinjection in the center of the LGd produces anterogradely-labeled fibers and retrogradely-labeled neurons along the rostrocaudal VISp that are located in a relatively central mediolateral position. Cases SW141021-01A and SW141021-02A both contain coinjection sites in the ventromedial half of the LGd but do not spread into the ventral strip area of LGd that borders the IGL. Both of these cases produced anterograde and retrograde labeling in the medial VISp adjacent to the border with VISam/VISpm. ARA, Allen Reference Atlas; BDA, biotinynlated dextran amine; CTB, cholera toxin subunit B; FG, Fluorogold; IGL, intergeniculate leaflet; LGd, dorsal lateral geniculate thalamic nucleus; LP, lateral posterior thalamic nucleus; PHAL, *phaseolus vulgaris* leucoagglutinin; VISam, anteromedial visual cortex; VISal, anterolateral visual cortex, VISl, lateral visual cortex, VISp, primary visual cortex; VISpl, posterolateral visual cortex; VISpm, posteromedial visual cortex.

**Figure 9.**
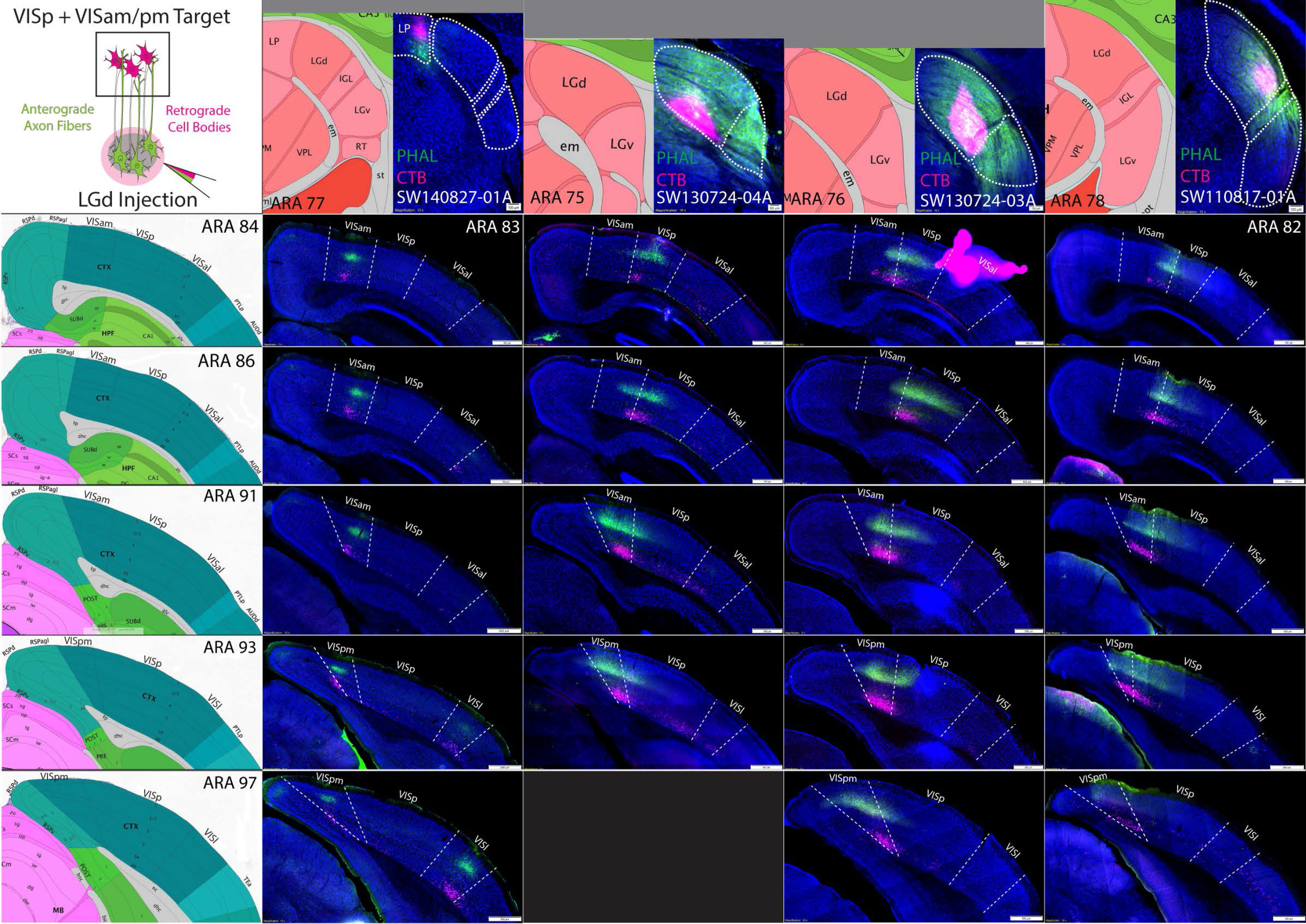
Anterograde and retrograde labeling pattern within VISp, VISam, VISpm after coinjections into the LGd that are located within the ventral strip region. A coinjection into the caudolateral LP near the border of the LGd and three coinjections into the ventral strip region of the LGd are shown (columns). The caudolateral LP coinjection produced anterogradely-labeled fibers and retrogradely-labeled cell bodies specifically within the VISam/pm that clearly identifies the VISam/pm boundary with VISp as demarcated by the representative ARA sections (rows). In comparison, the three LGd ventral strip coinjections have tracer labeling within the VISp that also crosses the boundary into the VISam and/or VISpm at multiple rostrocaudal levels. In all cases, retrogradely-labeled neurons are distributed within layer 6 while anterogradely-labeled fibers are distributed in layers 1, 4 and 6 with the greatest density in layer 4. Tissues sections that are not available at the relevant ARA level are omitted with a black square. ARA, Allen Reference Atlas; CTB, cholera toxin subunit B; IGL, intergeniculate leaflet; LGd, dorsal lateral geniculate thalamic nucleus; LP, lateral posterior thalamic nucleus; PHAL, *phaseolus vulgaris* leucoagglutinin; VISam, anteromedial visual cortex; VISal, anterolateral visual cortex, VISl, lateral visual cortex, VISp, primary visual cortex; VISpl, posterolateral visual cortex; VISpm, posteromedial visual cortex.

To confirm the VISam and VISpm boundaries with the extent of layer 4, we analyzed the Nissl cytoarchitecture in our tracer-labeled tissue sections and again found strong agreement of the distribution of tracer labeling within the VISam and VISpm (**Fig. 10**). After both LGd ventral strip coinjection (SW130724-04A) and caudal medial LP coinjection (SW140827-01A), PHAL-labeled fiber distribution is centered in layer 4 throughout the VISam and VISpm to the medial end of layer 4 where it meets the RSP.

**Figure 10.**
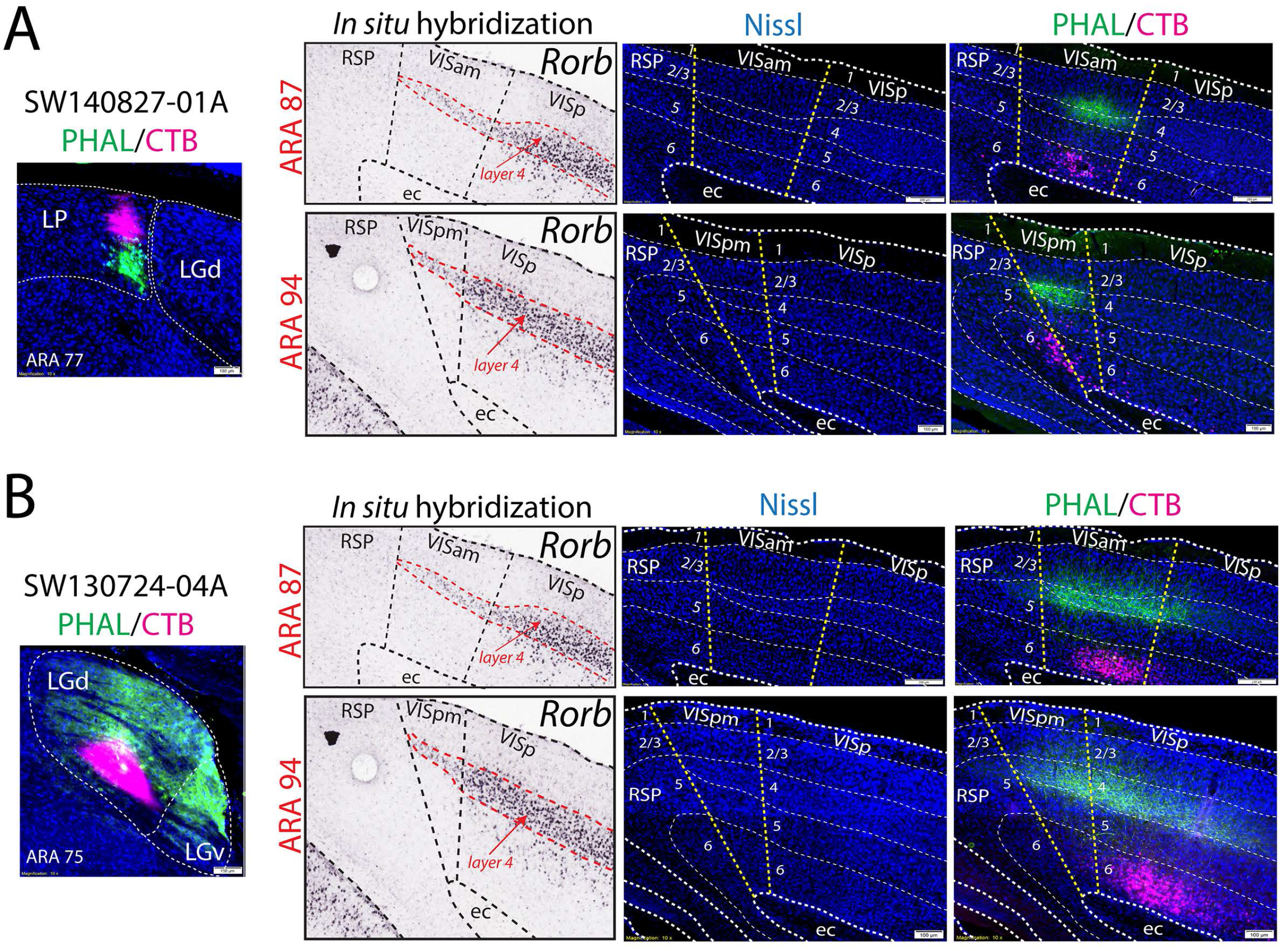
Anterograde and retrograde labeling patterns located within VISam and VISpm as defined by Nissl cytoarchitecture. The thin layer 4 of VISam and VISpm, as well as other cortical layers, can be identified by Nissl cytoarchitecture within our tracer-labeled tissue sections (injection sites shown on far left, *Rorb* gene expression on left, Nissl-only shown in the middle, Nissl plus tracer labeling on the right). At ARA levels 88 and 93, tracer labeling from an LP (SW140827-01A; **A**) and ventral strip LGd coinjection (SW130724-04A, **B**) are both localized within the VISam and VISpm. Anterogradely-labeled LGd fibers densely innervate layer 4 whereas robust numbers of retrogradely-labeled cortico-thalamic neurons are distributed within layer 6. ARA, Allen Reference Atlas; BDA, biotinynlated dextran amine; CTB, cholera toxin subunit B; ec, external capsule; FG, Fluorogold; IGL, intergeniculate leaflet; LGd, dorsal lateral geniculate thalamic nucleus; LP, lateral posterior thalamic nucleus; PHAL, *phaseolus vulgaris* leucoagglutinin; *Rorb,* RAR-related orphan receptor beta; RSP, retrosplenial cortex; VISam, anteromedial visual cortex; VISp, primary visual cortex; VISpm, posteromedial visual cortex.

### Distribution of tracer labeling within VISal, VISl, and VISpl after LGd coinjections

Following LGd coinjections, anterograde and retrograde labeling was observed in VISal, VISl, and VISpl (**Fig. 11A**). Regardless of coinjection site location within LGd, a sparse anterogradely-labeled terminal fields were distributed throughout VISal, VISl, and VISpl layer 4 whereas more robust numbers of retrogradely-labeled neurons were in VISal, VISl, and VISpl layer 6. In contrast to the presence of anterograde labeling in the lateral extrastriate areas after LGd coinjections, lateral extrastriate coinjections produced very few retrogradely-labeled neurons **(Fig. 11B)**. The rare few lateral extrastriate-projecting LGd neurons appeared scattered and inconsistent with the location of the descending fiber termination. Consistent with descending lateral extrastriate projections to LGd, tracer coinjection into different levels of the VISal or VISl anterogradely-labeled terminal fields in subregions of the LGd. Descending lateral extrastriate projections to the LGd were not as discrete as VISp projections, but in total, lateral extrastriate projections appear to innervate the entire LGd, including the VISam/VISpm-projecting ventral strip region (**Fig. 12**). In case SW111004-02A, double coinjection of BDA/FG into the VISam and PHAL/CTB into the VISal revealed that, in addition to direction connections between the VISam and VISal, VISal anterogradely-labeled fibers overlap with retrogradely-labeled VISam-projecting LGd neurons, suggesting a subcortical circuit between the medial and lateral extrastriate areas via the LGd ventral strip region (**Fig. 11c**).

**Figure 11.**
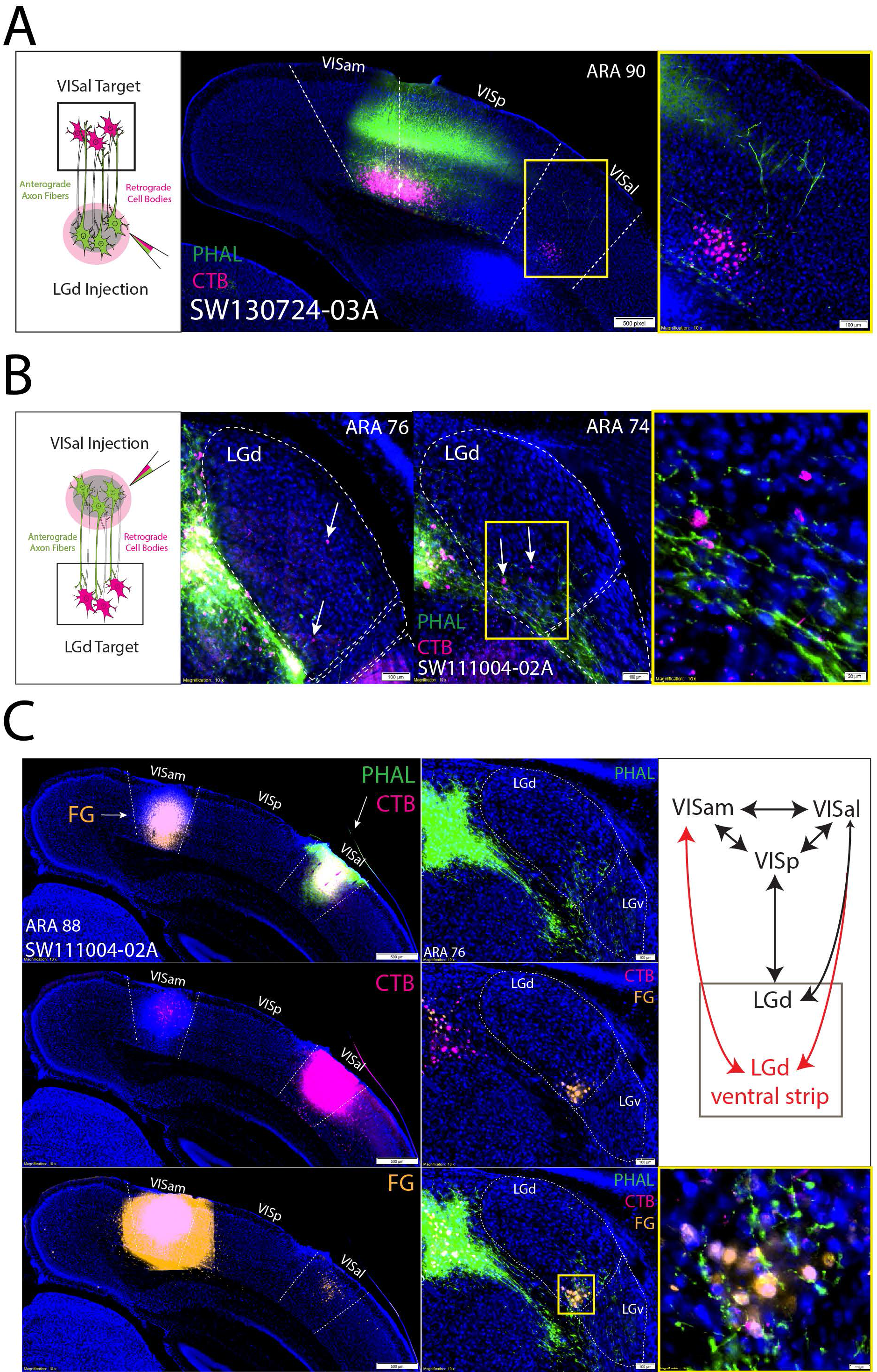
LGd connections with lateral extrastriate areas. (**A**) Anterograde and retrograde coinjection into all parts of the LGd produced sparse layer 4 anterograde labeling and robust layer 6 retrograde labeling in the lateral extrastriate visual areas. In SW130724-03A, PHAL/CTB coinjection into the LGd ventral strip region (injection site shown in **Fig. 8**) produced anterograde and retrograde labeling in the VISal that is segregated from the more dense terminal field in VISp/VISam (image on right is magnified view of yellow rectangle). This anterograde and retrograde labeling pattern was consistently observed in all LGd coinjection cases and within VISal, VISl, and VISpl. (**B**) Coinjections into the lateral extrastriate areas produced labeling within LGd that confirms the LGd coinjection data. In SW111004-02A, PHAL/CTB coinjection into the VISal reveals anterogradely-labeled PHAL terminal fields (green) and a few scattered retrogradely-labeled CTB neurons (magenta, demarcated by arrows; image on right is magnified view of yellow rectangle). For additional examples of lateral extrastriate coinjection tracer labeling within rostrocaudal LGd levels, see **Fig. 12**. (**C**) In case SW1110004-02A, we also coinjected BDA/FG into the VISam (BDA labeling too weak) in addition to PHAL/CTB in the VISam (top left). The accuracy of the two coinjection sites was clear as CTB injection into the VISal produced retrograde labeling in VISam layer 2/3 which overlapped the FG injection site (middle left) and FG injection in the VISam produced retrograde labeling in VISal layer 2/3 that overlapped the CTB injection site (bottom left). Note the injection sites in these images appear larger due to oversaturation necessary to visualize the dimmer retrograde labeling in the same section. Within the LGd of SW111004-02A, PHAL-labeled VISal terminal fields overlapped with retrogradely-labeled VISam-projecting LGd neurons (middle column, image on bottom right is magnified view of yellow rectangle). Overall, this data suggests a novel pathway for communication between the medial and lateral extrastriate visual areas via a subcortical pathway (red) through the LGd ventral strip (top right diagram). ARA, Allen Reference Atlas; BDA, biotinynlated dextran amine; CTB, cholera toxin subunit B; FG, Fluorogold; IGL, intergeniculate leaflet; LGd, dorsal lateral geniculate thalamic nucleus; LP, lateral posterior thalamic nucleus; PHAL, *phaseolus vulgaris* leucoagglutinin; VISam, anteromedial visual cortex; VISal, anterolateral visual cortex, VISl, lateral visual cortex, VISp, primary visual cortex.

**Figure 12.**
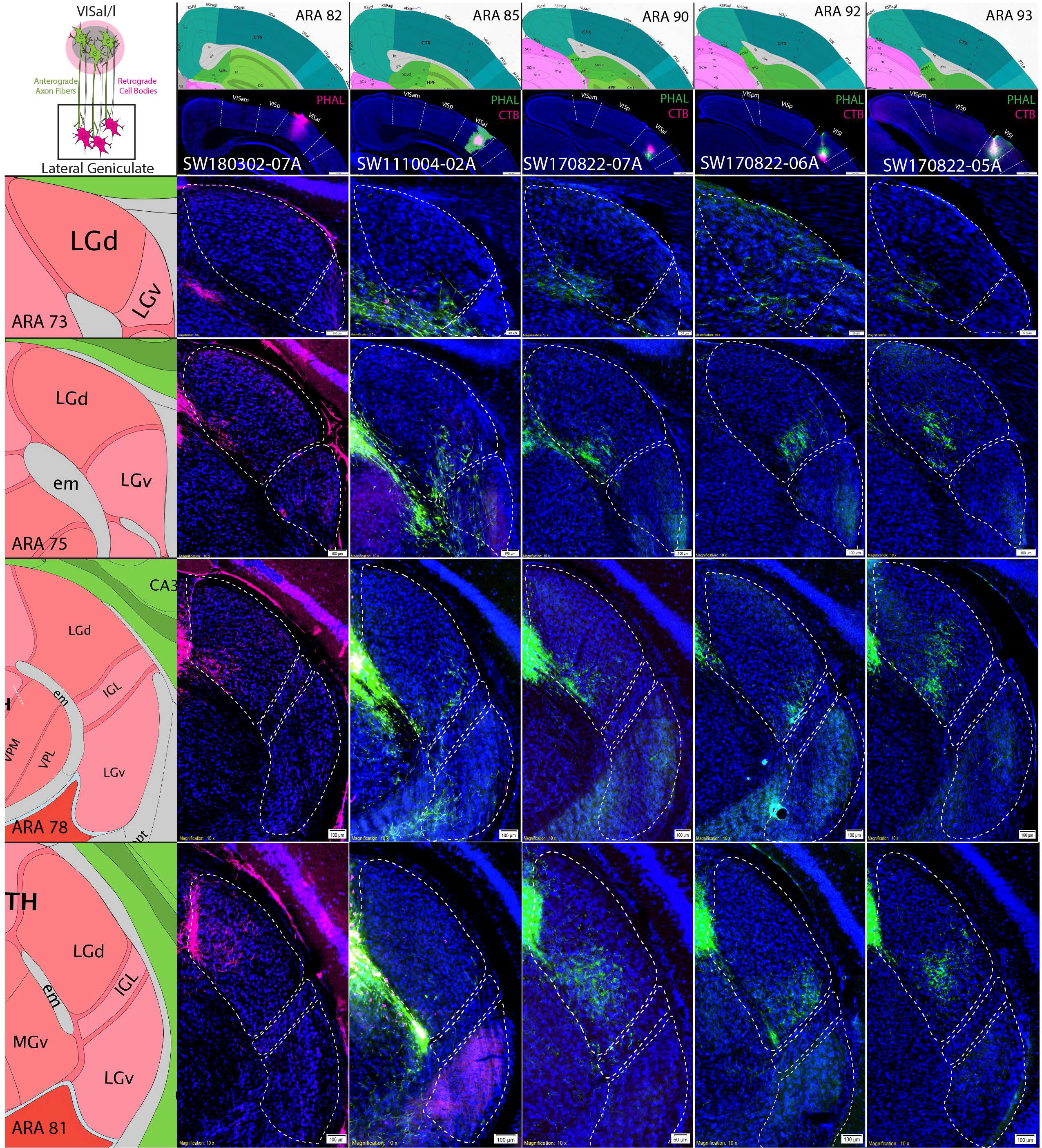
Distribution of tracer labeling in LGd after coinjection into the lateral extrastriate areas. Five coinjections of anterograde/retrograde tracer into lateral extrastriate areas at different rostrocaudal levels (columns) are shown with their labeling patterns in different rostrocaudal LGd levels (rows). For each column, extrastriate area tracer coinjection site is shown at the top with adjacent ARA atlas level (VISal = SW180302-07A, SW111004-02A, SW170822-07A; VISl = SW170822-06A and SW170822-05A). For each row across, the distribution of labeling throughout the rostrocaudal LGd is shown (representative ARA level shown at left). Each experimental case shows relatively discrete anterogradely-labeled PHAL terminal fields in different parts of LGd across multiple rostrocaudal levels. A few retrogradely-labeled LGd neurons were observed in SW111004-02A. ARA, Allen Reference Atlas; CTB, cholera toxin subunit B; IGL, intergeniculate leaflet; LGd, dorsal lateral geniculate thalamic nucleus; PHAL, *phaseolus vulgaris* leucoagglutinin; VISam, anteromedial visual cortex; VISal, anterolateral visual cortex, VISl, lateral visual cortex, VISp, primary visual cortex; VISpm, posteromedial visual cortex.

## Discussion

The results of this study support the existence of LGd extrastriate projections in the mouse and further extend on these findings to describe the overall thalamocortical organization of the LGd (**Fig. 13**). We have confirmed the previous report of LGd output to VISal and VISl as well as LGd input from VISam, VISal, and VISl (but not from the temporal association area) (Oh et al., 2014). In addition, we found evidence for LGd output to the VISam and bidirectional connectivity with the VISpm and VISpl. Our bidirectional circuit tracing strategy cross-validates the LGd extrastriate projections through localization of anterogradely-labeled LGd fibers within extrastriate regions and retrogradely-labeled extrastriate-projecting LGd neurons. Through this method, we characterized VIS cortex/LGd bidirectional connections both by laminar cell body location and projection fiber distribution. Overall, we provide evidence that neurons within a ventral LGd subregion provide bidirectional topographic connections with the medial extrastriate regions. In contrast, lateral extrastriate regions receive sparse non-topographic input from LGd but provide robust descending input to the ventral LGd region, including the medial extrastriate-projecting LGd neurons in the LGd ventral strip. Altogether, this reveals a potential disynaptic pathway of VISal→LGd→VISam, which suggests that the LGd may serve as interface for a unidirectional interaction of two extrastriatal areas (VISal and VISam), although functional significance of this pathway remain to be clarified.

**Figure 13.**
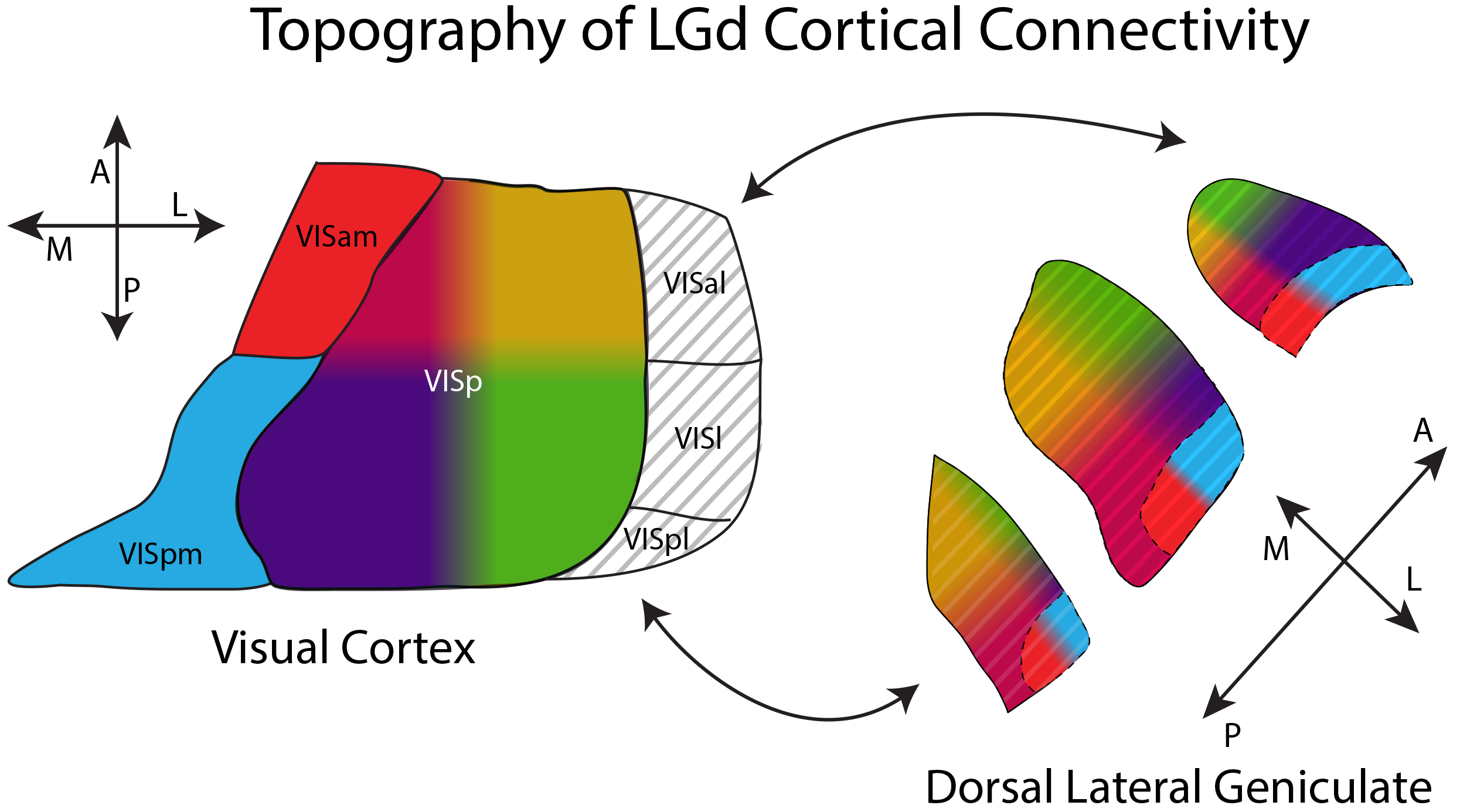
Topographic Organization of Mouse LGd Cortical Connectivity. The four quadrants of the VISp (see directional cross at bottom right; anteromedial (dark red), posteromedial (dark blue), anterolateral (yellow), posterolateral (green)) are bidirectionally-connected with LGd neurons in corresponding regions on the left (three LGd atlas level shown, related to directional cross at left). Connections of the VISam (red) and VISpm (blue) with the ventral strip region of LGd are a natural extension of the LGd/VISp topography. In contrast, VISal/l/pl connections with the LGd are primarily corticothalamic and terminate within LGd subregions (including the ventral strip region). Ascending LGd connections with the VISal/l/pl are likely scattered intermittently throughout the LGd and are not shown. A, anterior; L, lateral; LGd, dorsal lateral geniculate thalamic nucleus; M, medial; P, posterior; VISam, anteromedial visual cortex; VISal, anterolateral visual cortex, VISl, lateral visual cortex, VISp, primary visual cortex; VISpl, posterolateral visual cortex; VISpm, posteromedial visual cortex.

Our finding of an extrastriate-projecting ventral strip region adds novel understanding to the organization of the rodent LGd which can be divided into a core and superficial shell region and contains three morphologically-and electrophysiologically-distinct cell types (‘X-like’, ‘Y-like’, and ‘W-like’) for parallel visual processing(Krahe, El-Danaf, Dilger, Henderson, & Guido, 2011; Reese, 1988). Notably, LGd neurons with ‘X-like’ morphological and electrophysiological characteristics were shown to be strongly distributed in an area that coincides with the VISam/VISpm-projecting LGd ventral strip region(Krahe et al., 2011). In contrast, the superficial LGd shell region contains higher numbers of ‘W-like’ cells and is distinctly identified from the core region by input from the superior colliculus and direction-selective retinal ganglion cells (Bickford, Zhou, Krahe, Govindaiah, & Guido, 2015; Krahe et al., 2011; Reese, 1988). Recently, LGd neurons in the superficial shell region have been shown to themselves be anterior or posterior direction-selective allowing for horizontal-axis motion selectivity(Marshel, Kaye, Nauhaus, & Callaway, 2012). Interestingly, the most lateral part of the ventral strip LGd region containing VISpm-projecting neurons appears to intersect with the superficial shell region, suggesting that VISpm could receive input from both ‘X-like’ and ‘W-like’ LGd neurons (also superior collicular and direction-selective visual information), whereas VISam may only receive input from ‘X-like’ LGd neurons.

### LGd connections with the VISp

The results of our study are consistent with previous reports of bidirectional topographic connections between the LGd and VISp. LGd axon fibers primarily target VISp layers 1, 4, and 6 and LGd neurons receive input from VISp layer 6 neurons within defined topographic areas. The size and spherical shape of labeling clusters in LGd after VISp injection is likely a reflection of the size and shape of the coinjection site. Interestingly, coinjection sites that were located along the medial border of the VISp did not produce labeling within the ventral strip region of the LGd adjacent to the IGL border. Labeling in this ventral strip area was found only after coinjection sites that were centered in the VISam or VISpm.

### Extrastriate LGd connections with the VISam and VISpm

Coinjections of anterograde and retrograde tracer into the VISam and VISpm produced clustered labeling within the ventral strip region of the LGd in a topographic manner. One possible interpretation of this result could be that the coinjection sites in VISam and VISpm spread into the laterally adjacent medial VISp area and LGd labeling in the ventral strip region is actually from VISp. However, our data suggests multiple points of evidence to the contrary. First, coinjections into the medial VISp do not produce labeling within the ventral strip LGd region. Second, coinjection sites which are located near the medial RSP border of the VISam/VISpm (away from the VISp) produced tracer labeling within the LGd ventral strip. Third, LGd coinjection sites that are centered in or spread into the LGd ventral strip region produce anterograde and retrograde labeling within the VISam and VISpm. Fourth, LGd coinjection sites that are located near to, but do not spread into, the LGd ventral strip region produced anterograde and retrograde labeling within the medial VISp that is immediately adjacent to the VISam/VISpm border.

Another possible interpretation could be that the boundaries of the VISam and VISpm are not accurate and the VISp extends further medially then shown in the ARA. Neuroanatomical atlases provide good approximations, but oblique histological tissue sectioning can produce incongruencies when comparing experimental data to atlas sections. Our analysis of gene expression, cytoarchitecture, and other connectivity patterns all confirmed the accuracy of the VISp and extrastriate area boundaries as shown in the ARA.

### Extrastriate connections of the LGd with the VISal, VISl, and VISpl

In all our LGd coinjection cases, we observed sparse anterogradely-labeled fibers in layer 4 and relatively more dense numbers of retrogradely-labeled neurons in layer 6 in the lateral extrastriate regions. Lateral extrastriate coinjections confirmed the descending lateral extrastriate axon fibers with each coinjection terminating in different subregions of LGd. Notably the distribution of VISal and VISl fibers within the ventral parts of the LGd is entirely different from the adjacent lateral VISp whose fibers target the dorsal parts of LGd. Therefore, our VISal and VISl anterograde labeling cannot be the result of coinjection site spread into the VISp. In contrast, the ascending LGd projection to lateral extrastriate areas was more difficult to verify as very few retrogradely-labeled neurons were present in the LGd after lateral extrastriate coinjections and the distribution of those rare cells appeared to be randomly scattered. One possibility could be that the fiber labeling is so sparse and the coinjection sites are so small, that only a few cells project within the area of retrograde tracer uptake. In support of this, many of the studies that reported positive findings of extrastriate LGd projections in the larger non-human primate brain reported that retrograde labeling from lateral extrastriate regions was ‘sparse’ and ‘scattered’ with one non-human primate study reporting retrogradely-labeled neurons numbering ~20 neurons per section (**Table 1**). To test this, we performed a quadruple retrograde injection experiment along the rostrocaudal axis of the lateral extrastriate regions to maximize the likelihood of retrograde labeling (data not shown). The quadruple retrograde tracing approach did manage to retrogradely-label more neurons, but the total number of retrogradely-labeled neurons was still underwhelming (<5 total neurons). Overall, our interpretation is that there is a minor scattered subpopulation of LGd neurons that project to lateral extrastriate visual areas. In turn, the lateral extrastriate visual areas send dense descending projections to topographic parts of LGd, including the ventral strip, suggesting a subcortical LGd pathway for lateral extrastriate influence on medial extrastriate regions.

### Conclusions

The existence of extrastriate LGd connectivity has been debated for decades in a variety of different animal models. We provide evidence that the mouse LGd had extensive connections with both the medial and lateral extrastriate regions. VISam-and VISpm-projecting LGd neurons are located along a ventral strip region adjacent to the IGL border. These findings provide new insight into the organization of the rodent LGd and its thalamocortical connections.

